# Phytoplankton size structure and biogeochemical responses to nutrient enrichment in an oligotrophic coral reef

**DOI:** 10.64898/2026.04.06.716629

**Authors:** Jorge L. Suarez-Caballero, Takashi Nakamura

## Abstract

Tropical coral reef ecosystems worldwide are being impacted by combined pressures of climate change and human activities that introduce large quantities of nutrients and sediments into coastal areas. In this context, phytoplankton represent a critical link between dissolved inorganic nutrients and coral reef food webs, yet their role in these ecosystems remains understudied. We investigated ecological responses of the summer phytoplankton community of Shiraho Reef (Ishigaki Island, Okinawa, Japan) to nutrient enrichment using field-based microcosm experiments under natural light and temperature conditions in September 2022 and 2023. Treatments included single and combined additions of nitrogen, phosphorus, and silicon. Chlorophyll *a* (Chl *a*) concentrations increased after three days under combined nutrient conditions, whereas single-nutrient additions produced limited responses, indicating a strong co-limitation by nitrogen and phosphorus in the reef. Analysis of size-fractionated Chl *a* revealed shifts from picophytoplankton that typically dominate tropical oligotrophic ecosystems toward larger groups supported by enhanced nutrient availability. Our results show short-term impacts of nutrient enrichment events on phytoplankton size structure and biogeochemical cycling in coral reefs, and highlight the importance of pelagic processes in coral reef carbon dynamics under nutrient-enrichment.

## Introduction

Tropical coral reefs are among the most diverse and productive ecosystems on Earth (Odum and Odum 1955; Connell 1978; Allgeier 2024). Despite nutrient-poor conditions, these ecosystems support high primary productivity and complex trophic webs, commonly attributed to efficient nutrient recycling and tight biological coupling between reef organisms (Nelson et al. 2023). Local and context-dependent environmental conditions, including hydrography, trophic status, seasonality, and extreme weather events, further influence the reef and water quality in the system (Delesalle et al. 1993; Tsuchiya et al. 2014; Page et al. 2023). Dissolved nutrients regulate primary productivity, and nitrogen (N) and phosphorus (P) are generally considered the major limiting nutrients in marine and freshwater environments, respectively (Hecky et al. 1988; Howarth and Marino 2006). However, numerous studies have found that synergistic limitation by multiple nutrients is common in aquatic ecosystems, especially involving nitrogen and phosphorus (Elser et al. 2007; Harpole et al. 2011). Such interactions may be strongly influenced by local conditions, including water residence time, habitat type, nutrient stoichiometry, and nutrient inputs (Downing et al. 1999). Coral reefs are characterized by clear water and low dissolved nutrient concentrations. However, coastal systems are increasingly affected by human activities that enhance nutrient inputs through river runoff, sewage, and groundwater discharge, which can lead to changes in marine water chemistry and ecosystem metabolism (Koop et al. 2001; Furnas et al. 2005; Silbiger et al. 2025). Although ecological consequences of eutrophication have been recognized for decades (Ryther *et al*. 1971; Vollenweider 1992), massive inputs of nutrients from land use change, agriculture, and urbanization are altering global nutrient cycles and causing severe adverse effects on coastal ecosystems (Conley et al. 2009; Richardson et al. 2023). These include coral reef degradation (Fabricius 2005; D’Angelo and Wiedenmann 2014), harmful algal blooms (HABs) (Anderson et al. 2002; Heisler et al. 2008), and community changes with direct and indirect cascading effects on food web dynamics and water biogeochemistry (Smith et al. 1999; Wurtsbaugh et al. 2019; Silbiger et al. 2025).

Research focusing on these ecosystems has traditionally emphasized benthic organisms, i.e., corals and macroalgae, as the main drivers of primary production, calcification, and nutrient cycling, often treating pelagic processes as secondary or negligible components. However, recent studies have emphasized the importance of microbial processes in the water column, especially under nutrient enrichment in shallow coastal areas (Nelson et al. 2023; Shakya and Allgeier 2023). Phytoplankton communities form the base of marine food webs, linking inorganic compounds to biological systems via photosynthesis, transferring matter and energy to higher trophic levels. Chlorophyll *a* (Chl *a*), the most ubiquitous pigment in phytoplankton, is commonly used as a proxy for their carbon biomass (Iriarte and Purdie 1994; Hirata et al. 2008). The oligotrophic nature of coral reefs typically favors small phytoplankton size classes with efficient nutrient uptake due to their high surface-to-volume ratios, whereas larger phytoplankton are at a competitive disadvantage under low-nutrient conditions (Marañón et al. 2015; Ward et al. 2017; Beltrán-Heredia et al. 2017). Therefore, picophytoplankton often dominate total Chl *a* concentrations in low-biomass tropical reef waters, whereas nano- and microphytoplankton contribute relatively less under oligotrophic conditions (Furnas and Mitchell 1987; Iriarte and Purdie 1994; Tada et al. 2004; Wei et al. 2022). In contrast, nutrient-enriched environments with higher chlorophyll concentrations support larger phytoplankton groups (Iriarte and Purdie 1994; Yi et al. 2014).

Experimental approaches employing microcosms and mesocosms have frequently been used to study phytoplankton responses to specific environmental factors, providing essential information on how they influence primary productivity (Sommer 1988; Schlüter 1998; Yoon et al. 2024). Although extensive research has been conducted in temperate regions, upwelling zones with high seasonal productivity, and subtropical gyres, phytoplankton dynamics in tropical coral reefs have received less attention. In this study, we investigate nutrient-driven changes in phytoplankton size structure and water chemistry in a fringing coral reef at Ishigaki Island, Okinawa, Japan. Shiraho Reef has a long history of research dating back more than three decades, with studies documenting the hydrodynamics, benthic structure, carbon fluxes, and community metabolism (Nakamori et al. 1992; Kayanne et al. 1995; Hata et al. 2002). Subsequent studies focused on the importance of terrestrial nutrient inputs and other processes influencing reef biogeochemistry (Umezawa et al. 2002; Blanco et al. 2010; Sith et al. 2019). Despite establishing Shiraho as a model system for understanding terrestrial influences and benthic processes, biological dynamics in the water column have received considerably less attention. To address this gap, we conducted field-based microcosm experiments during summer, when nutrient enrichment occurs due to higher rainfall, groundwater discharge, and typhoons (Blanco et al. 2010). By analyzing Chl *a* in size-fractionated phytoplankton samples with specific nutrient additions (N, P, Si), we sought to clarify short-term impacts of nutrient enrichment events in coral reefs exposed to terrestrial anthropogenic pressures. We hypothesized that nutrient enrichment at coral reefs can rapidly alter the structure of phytoplankton communities and influence biogeochemical cycling, including carbon sequestration, and nutrient limitation patterns.

Understanding these contributions of planktonic communities in coral reefs is necessary to accurately quantify primary productivity, carbon dynamics, and ecosystem responses, especially under climate change and anthropogenic pressures in coastal areas.

## Materials and Methods

### Study site

Ishigaki Island (24º30’N, 124º10’E) is a subtropical island in the southwestern Ryukyu Archipelago, Japan (Fig. 1a). It is influenced by the Kuroshio Current, which transports warm, oligotrophic water from the tropics and strongly influences the local marine environment (Liu et al. 2021; Durán Gómez and Nagai 2022). The island is surrounded by extensive fringing coral reefs that protect the shoreline from wave action. On the southeastern side, Shiraho Reef (24° 21’ 52.47” N, 124° 15’ 12.25” E) is a semi-enclosed reef that extends several hundred meters from the shore, with a reef flat (∼850 m) leading to the reef crest (Fig. 1b-c). Several seagrass patches, coral, and algal turf, contribute to its high productivity (Kayanne et al. 1995). Water circulation and mixing in the reef are primarily driven by tidal currents, wind, and waves, involving an inflow of oceanic water into the backreef moat and outflow through channels back to the open sea (Nakamori et al. 1992). Additionally, the reef is influenced by terrestrial inputs from adjacent rivers, particularly the Todoroki River, which delivers significant nutrients and sediment to the coastal lagoon, especially in summer (Kawahata 2000; Hata et al. 2002; Blanco et al. 2010; Sith et al. 2017).

**Figure 1.**
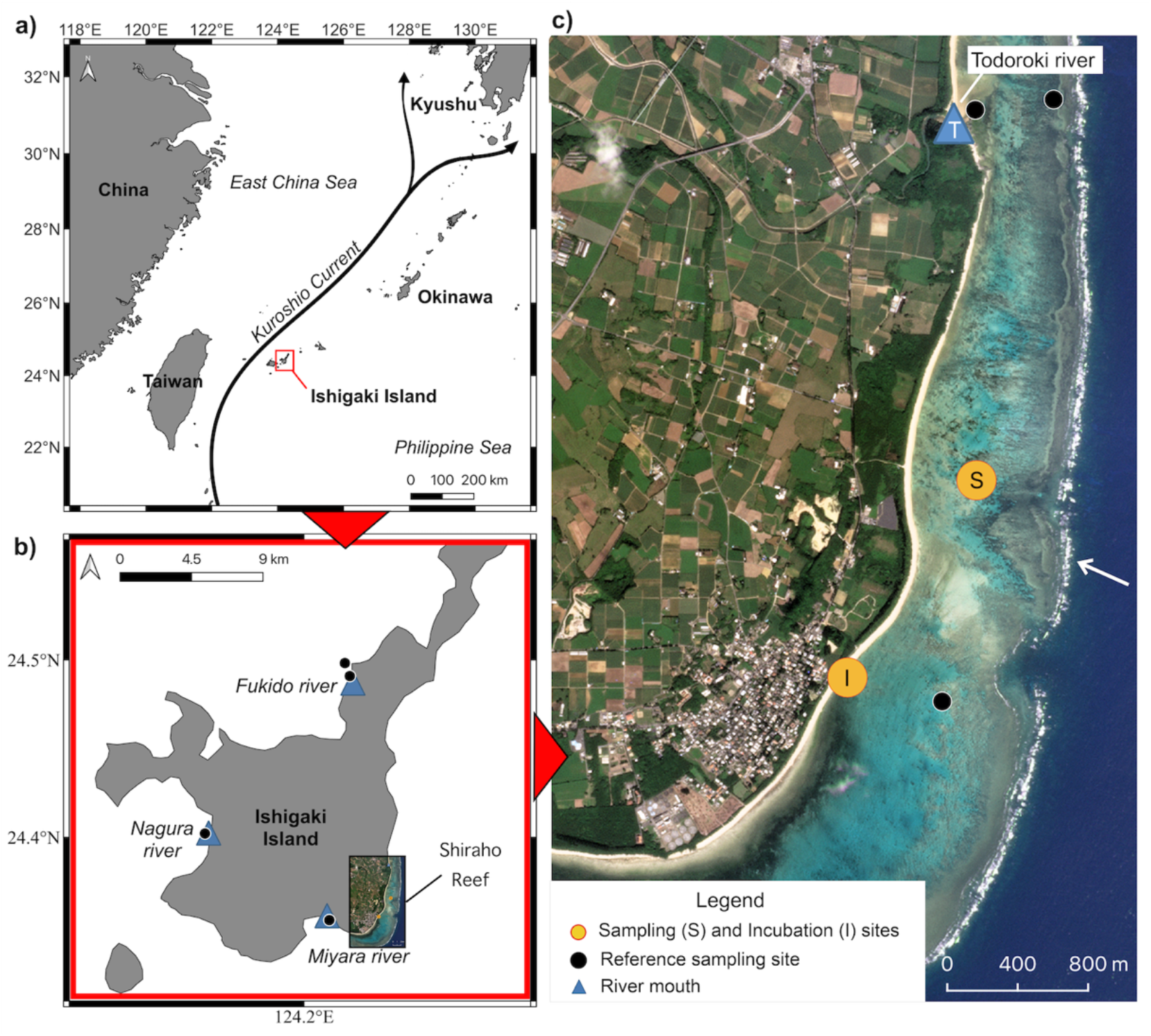
Map of the study site: (a) location of Ishigaki Island (red box) and the Kuroshio Current, (b) Ishigaki Island showing Shiraho Reef and additional sampling sites around the island (black points) including nearby river mouths (blue triangles), (c) satellite view of the southeastern area showing the coral reef crest (white arrow), microcosm sampling site (S), incubation site at Shiraho Beach (I), and Todoroki River mouth (blue triangle) Satellite image: Planet Labs PBC (2025).

### Microcosm experiments

Microcosm experiments were conducted twice for seven consecutive days in September 2022 and 2023 with natural phytoplankton communities. Approximately 150 L of surface seawater were collected from the Shiraho Reef area by swimming, using three clean 50 L polyethylene (PE) containers. Each one was treated as an independent experimental replicate. Seawater samples were pre-screened through a 150-µm nylon mesh to exclude large debris and zooplankton that could negatively impact primary production due to grazing. After screening, seawater was distributed into 12 transparent 10-L PE bottles, corresponding to four treatments with three independent replicates, where nutrient additions were performed: (1) Control: no added nutrients, (2) N: 20 µM nitrate, (3) P: 1.25 µM phosphate, and (4) NP: 20 µM nitrate + 1.25 µM phosphate. In 2023, we introduced an additional treatment (5) NPSi: 20 µM nitrate + 1.25 µM phosphate + 20 µM silicate, to test effects without silicate limitation. To adjust concentrations, nutrient stock solutions were prepared with sodium nitrate (NaNO_3_), potassium phosphate (KH_2_PO_4_), and sodium hexafluorosilicate (Na_2_SiF_6_) and added with a micropipette to achieve the intended concentrations, simulating a pulse of nutrients in the environment. Enrichment was performed using an N:P ratio of 16:1, according to Redfield stoichiometry (Redfield 1958; Redfield et al. 1963), and a N:Si ratio of 1:1 in the NPSi treatment, based on mean diatom requirements (Brzezinski 1985). An additional control bottle was equipped with a Photosynthetically Active Radiation (PAR) light sensor (DEFI2-L, JFE Advantech Co., Ltd., Japan) and a temperature logger (HOBO, Onset, USA) to monitor these parameters in the containers. All bottles were kept underwater at Shiraho beach, attached with a net to the seafloor at 2 m depth to maintain natural temperature and light conditions in the field, while allowing wave movement to prevent sedimentation. Air was removed entirely from the bottles to prevent physical damage to phytoplankton by air bubbles.

### Subsampling and laboratory analyses

Initial conditions and daily subsamples were collected from each container to measure Chl *a*, dissolved inorganic nutrients, dissolved organic carbon (DOC) concentrations, and carbonate system parameters. For total Chl *a* extraction, approximately 300 mL of water were vacuum filtered onto GF/F glass microfiber filters (0.7 µm pore size, 25 mm diameter; Whatman, Cytiva, UK), extracted with N, N-dimethylformamide (DMF), and determined with a fluorometer (Turner Designs, Model 10-AU, USA) (Suzuki and Ishimaru, 1990). The exact filtered volume was recorded for each sample and used to calculate Chl *a* concentrations. In the 2023 experiment, Chl *a* was determined for three phytoplankton size fractions: microphytoplankton (10 – 150 µm), nanophytoplankton (1 – 10 µm), and picophytoplankton (0.2 – 1 µm). These fractions were obtained by sequential filtration through filters with pore sizes of 10 µm (GA-100, Advantec, Japan), 1 µm (PTFE Omnipore Merck Millipore, USA), and 0.2 µm (PFTE Advantec, Japan). The choice of filters was based on chemical compatibility with DFM, used for pigment extraction, to avoid degradation that might influence fluorescence measurements. Although these operational size fractions differ slightly from traditional conceptual phytoplankton size classifications, they are commonly applied in size-fractionated chlorophyll studies in oligotrophic sub-tropical oceans (Iriarte and Purdie 1994). During the subsampling routine, additional samples were taken simultaneously with GF/F filters to ensure consistency with the size fractionation. Agreement between the sum of size-fractionated and total Chl *a* extracted with GF/F filters supported the reliability of the size-fractionation method for quantifying Chl *a* (Figs. S1 and S2). For dissolved inorganic nutrient analysis, approximately 50 mL of sample were filtered through 0.45-µm acetate membrane filters (Advantec, Japan) into PE tubes and immediately frozen at -20 ºC until analysis. Concentrations of dissolved nitrate (NO_3_^-^), nitrite (NO_2_^-^), ammonium (NH_4_^+^), phosphate (PO_4_^3-^), and silicate (SiO_2_) were determined using a QuAAtro 2-HR flow auto-analyzer (SEAL Analytical Ltd., Germany; BLTEC, Japan) following manufacturer protocols and best-practice colorimetric methods for seawater analysis (Becker et al. 2020). For DOC analysis, 50 mL of seawater sample were filtered through pre-combusted GF/F filters (450°C for 4 h) using a glass syringe. Filtrate was collected in acid-washed, pre-combusted glass vials (10 mL), and stored at -20 ºC until analysis. DOC concentrations were quantified via high-temperature catalytic oxidation (Sugimura and Suzuki 1988) with a total organic carbon analyzer (TOC-L, Shimadzu Corporation, Japan). For carbonate system analysis, samples were collected in 250-mL borosilicate glass bottles and fixed with 100 µL of saturated mercuric chloride (HgCl_2_) to stop biological activity. Bottles were sealed and stored in the dark until analysis. Total alkalinity (TA) and pH on the total scale (pH_T_) were measured to calculate dissolved inorganic carbon (DIC; CO_2_ + HCO_3_^-^ + CO_3_^2-^) and other carbonate system parameters. TA was determined using the open-cell titration method with an automatic titrator (ATT-15, Kimoto, Japan) and a glass electrode (Thermo Fisher Scientific, USA) calibrated with Tris and AMP buffer solutions, following the *Guide to Best Practices for Ocean CO*_*2*_ *Measurements* (Dickson et al. 2007). DIC concentrations were calculated using CO2SYS v3.0 (Pierrot et al. 2021), incorporating TA, pH, temperature, salinity, and silicate and phosphate measurements. Carbonate system calculations used the following constants: K_1_ and K_2_ from Millero (2010), KHSO_4_ from Dickson (1990), KHF from Perez and Fraga (1987), total boron [B]_T_ from Lee et al. (2010), and the EOS-80 equation of state for seawater. Certified reference materials for oceanic CO2 measurements, provided by A. Dickson (Scripps Institution of Oceanography), were periodically measured to assess precision and accuracy. The standard deviation of repeated CRM measurements ranged from 0.006–0.010 for pH_T_ and 1.98–4.42 µmol kg^-1^ for TA; these values were used as input uncertainty for error propagation in CO2SYS.

### Statistical analyses and nutrient uptake kinetics

Chl *a* results were analyzed with a linear mixed-effects model (LMM) fitted to log-transformed Chl *a* concentrations to detect statistically significant differences between treatments over time while accounting for unbalanced data and variability among independent replicates. Size-fractionated Chl *a* concentrations were evaluated using Tukey-adjusted pairwise comparisons of estimated marginal means (EMMs) among treatment, day, and filter size combinations. Consistency between total and size-fractionated Chl *a* samples was assessed by linear regression and Spearman’s correlation analysis. For calculated seawater carbonate system parameters, error propagation was performed using the integrated uncertainty module in CO2SYS v3.0. All analyses were conducted using Python 3.12, with pandas, numpy, and scipy.stats for statistical analysis, and plotted using the matplotlib and seaborn libraries. LMMs were performed with R (R Core Team 2025) using the lme4 package (Bates et al. 2015). Nutrient uptake kinetics were modeled using a Hill-type function accounting for co-limitation by nitrogen, phosphorus, and silicate. Uptake of dissolved inorganic nitrogen (DIN; NO_2_^-^ + NO_3_^-^ + NH_4_^+^) was expressed as:

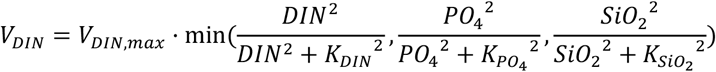

where *Vmax* is the maximum uptake rate and *K* is the half-saturation constant for each nutrient. The minimum operator reflects Liebig’s law of the minimum, assuming that uptake is constrained by the most limiting nutrient. The parameters *Vmax* and *K* for DIN, PO_4_^3-^, and SiO_2_ were estimated by nonlinear curve fitting. Model performance was evaluated using the Willmott Skill Index (Willmott 1981), and parameter values were iteratively optimized to maximize model performance across the three nutrient uptake rates. Analogous formulations for PO_4_^3-^ and SiO_2_ uptake, and the Willmott Skill Index formula are provided in the Supplementary Material (Table S1).

## Results

### Initial physicochemical parameters of seawater

Water temperature and light conditions fluctuated daily at the incubation site (Fig. 2). Mean daily temperatures in 2023 were consistently higher than in 2022 (30.98 ± 0.33 vs 28.3 ± 0.37 °C). Photon flux density (PFD) followed a ∼13:11-hour light-dark cycle in both years, with peak values reaching 2,651 μmol photons m^-2^ s^-1^ in 2022 and 1,993 μmol photons m^-2^ s^-1^ in 2023. Despite short-term variations, daily light integral did not differ significantly between years (Mann-Whitney U test, p > 0.5). Initial physicochemical parameters are summarized in Table 1. DIN and PO_4_^3-^ concentrations were higher in 2022 (2.56 and 0.23 µM) than in 2023 (0.8 and 0.07 µM), whereas SiO_2_ was similar in both years (2.63 and 2.13 µM). Initial Chl *a* concentration was higher in 2023 than in 2022 (0.65 vs. 0.55 µg L^-1^), while salinity, pH_T_, DOC, and TA concentrations showed only minor differences between years.

**Table 1.** Initial environmental parameters of microcosm experiments.

|  | September 11 – 17, | September 18 – 25, | Units |
| --- | --- | --- | --- |
|  | 2022 | 2023 |  |
| Salinity | $34.14 \pm 0.03$ | $34.12 \pm 0.01$ | psu |
| $\text{pH}_T$ | $8.14 \pm 0.001$ | $8.04 \pm 0.005$ | - |
| $\text{NO}_3^-$ | $1.18 \pm 0.06$ | $0.55^*$ | $\mu\text{M}$ |
| $\text{NO}_2^-$ | $0.15 \pm 0.02$ | $0.05^*$ | $\mu\text{M}$ |
| $\text{NH}_4^+$ | $1.24 \pm 0.25$ | $0.20^*$ | $\mu\text{M}$ |
| $\text{PO}_4^{3-}$ | $0.23 \pm 0.05$ | $0.07^*$ | $\mu\text{M}$ |
| **SiO<sub>2</sub>** | 2.63 ± 0.13 | 2.13* | μM |
| **DOC** | 76.19 ± 3.86 | 65.45 ± 3.37 | μM |
| **DIC** | 1915.1 ± 7.8 | 1986.4 ± 5.6 | μmol Kg<sup>-1</sup> |
| **TA** | 2262.83 ± 3.63 | 2278.58 ± 1.62 | μmol Kg<sup>-1</sup> |
| **Chl *a*** | 0.55 ± 0.06 | 0.65 ± 0.06 | μg L<sup>-1</sup> |
| **Chl *a* (0.2 – 1 μm)** | - | 5.9 ± 0.002 | % |
| **Chl *a* (1 – 10 μm)** | - | 16 ± 0.01 | % |
| **Chl *a* (10 – 150 μm)** | - | 78.1 ± 0.05 | % |

\*Only one measurement was taken. For all others, averages from independent replicates are shown (n=3).

**Figure 2.**
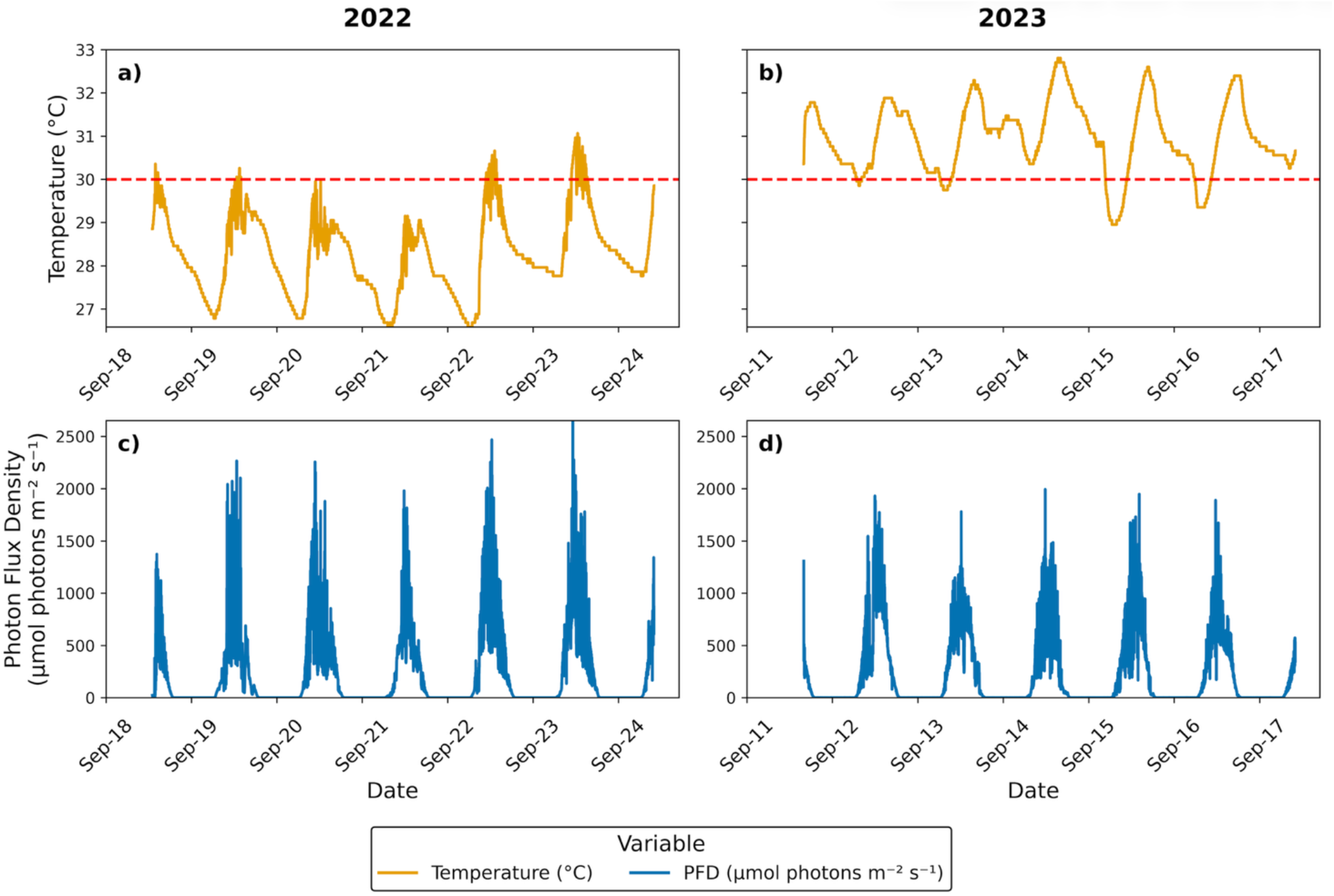
(a, b) Water temperature at the incubation site over 7-day experiments in 2022 and 2023. The horizontal dashed line represents 30 ºC. (c, d) Photon flux density (PFD) for the same periods.

### Dissolved inorganic nutrient dynamics

Dissolved inorganic nutrient concentrations before and after enrichment are summarized in Table 2. Throughout the 7-day incubation period, nutrient dynamics revealed clear treatment effects (Fig. 3). The control maintained low and stable DIN and PO_4_^3-^ concentrations. Single-nutrient treatments (N and P) showed limited changes, as their respective concentrations remained relatively constant throughout the experiment. In contrast, combined nutrient additions (NP) led to a rapid and sustained decline in both DIN and PO_4_^3-^, with an even stronger depletion observed in the NPSi treatment. Although DIN and PO_4_^3-^ were partially consumed during the first 48 h, SiO_2_ concentrations remained stable in all treatments during this initial phase. A sharp depletion of SiO_2_ was observed in the NPSi treatment of 2023 on Day 3, followed by a more gradual decline in the NP treatment on Day 4. Significant differences in SiO_2_ were observed only in N and NP treatments in 2022 and in NPSi in 2023. Despite differences in absolute concentrations between years, dissolved N:P ratios at Shiraho Reef were similar (11.13 and 11.43). While the NP and NPSi treatments exhibited nutrient ratios close to the Redfield ratio in both years, individual additions of P and N resulted in strongly imbalanced nutrient conditions. Complete nutrient data and stoichiometric ratios are available in the Supplementary Material (Figs. S3-S5; Table S2).

**Table 2.**
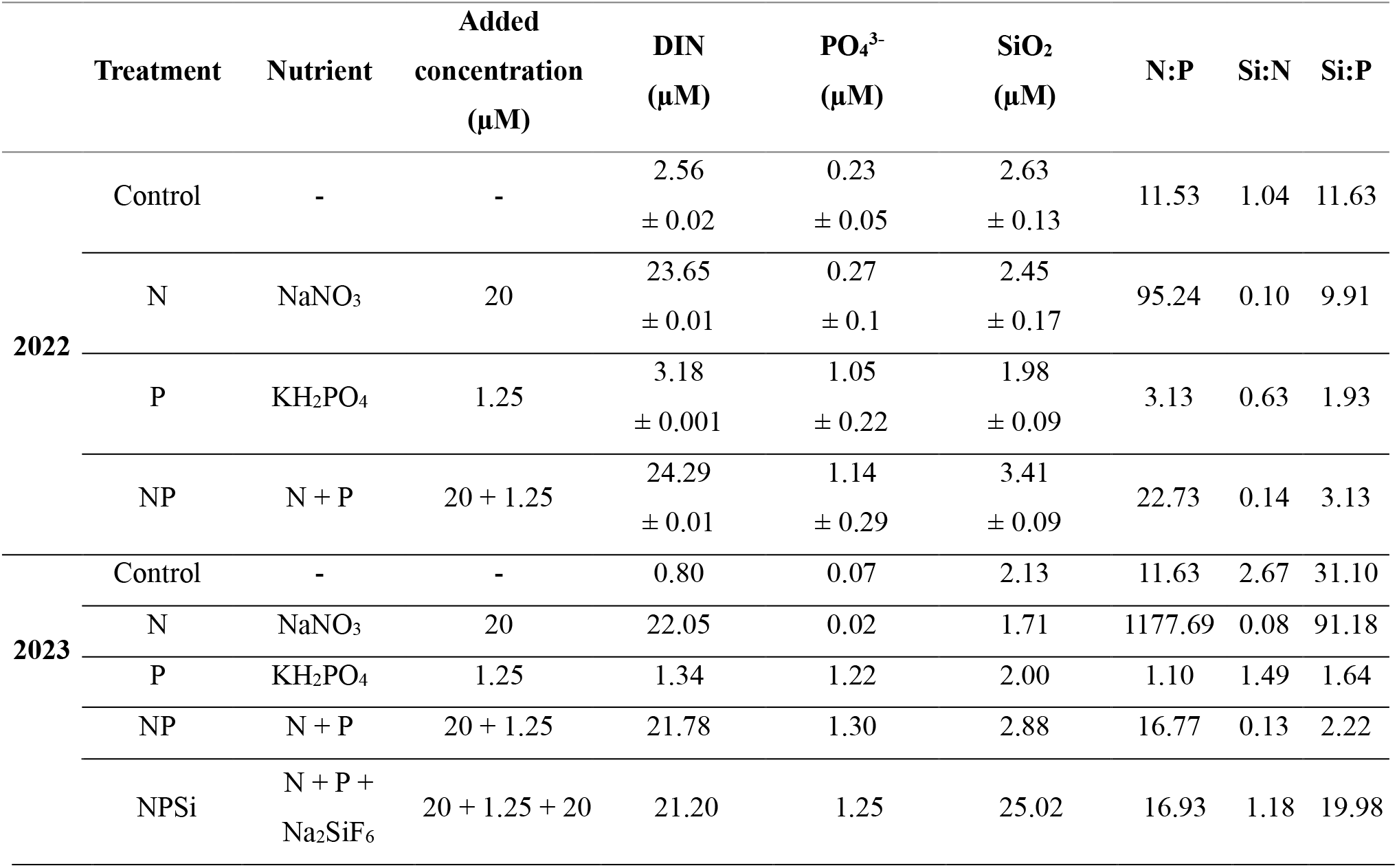
Dissolved inorganic nutrient concentrations before and after nutrient enrichment per treatment at the beginning of the experiment. N:P, Si:N, and Si:P values represent ratios of dissolved inorganic nutrients.

**Fig. 3.**
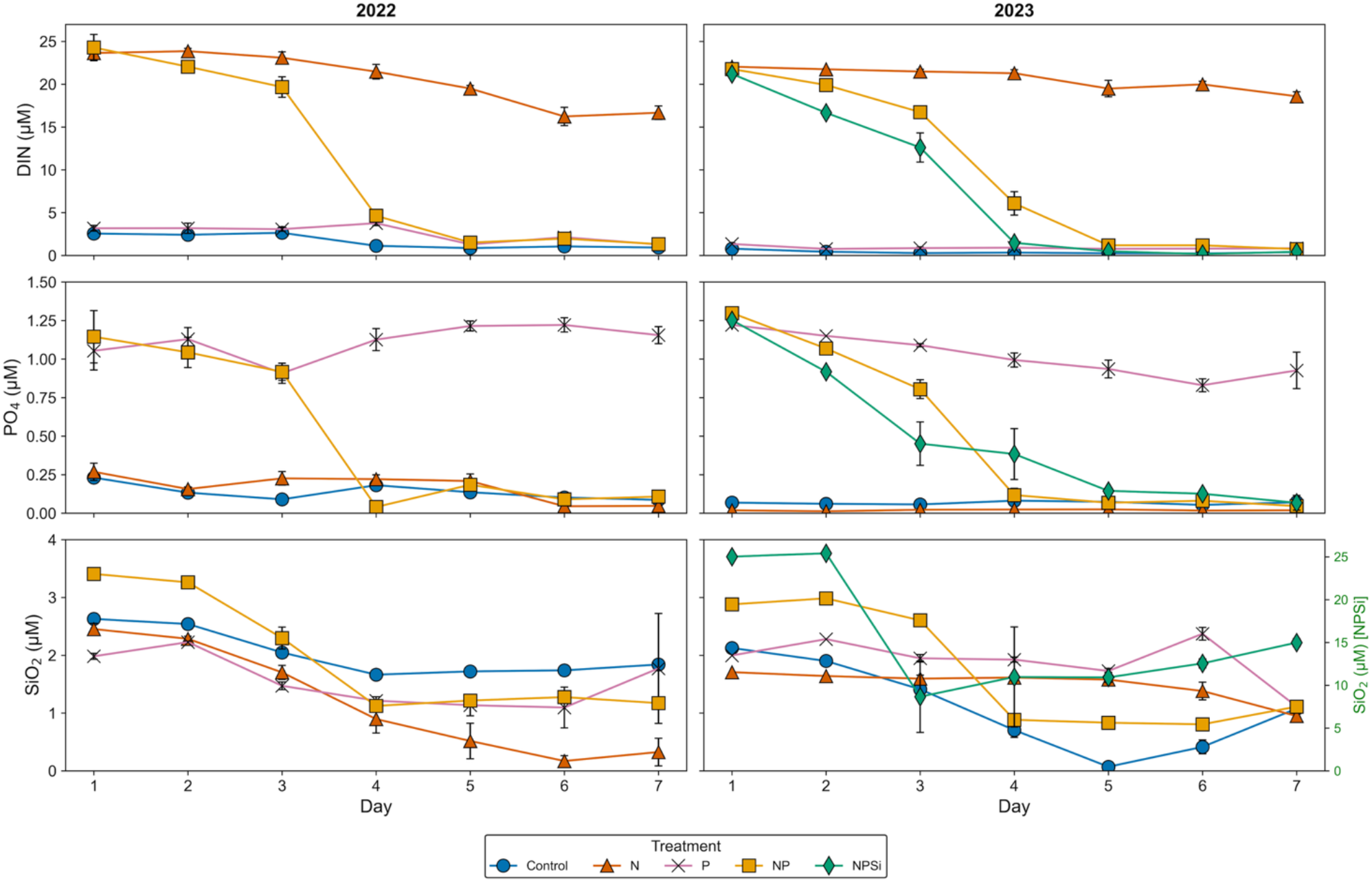
Temporal changes in dissolved inorganic nutrients over a 7-day incubation period in 2022 (left) and 2023 (right) under different nutrient treatments. Panels show dissolved inorganic nitrogen (DIN), dissolved phosphate (PO_4_^3-^), and dissolved silicate (SiO_2_). Colors indicate treatments: Control (blue), N (orange), P (pink), NP (yellow), and NPSi (green). The green secondary axis corresponds only to the NPSi treatment in 2023. Error bars represent standard error (±SE) calculated from independent replicates (n = 3).

### Nutrient uptake rates

Temporal dynamics of nutrient uptake revealed distinct responses to enrichment treatments (Fig. S6). NO_3_^-^ and PO_4_^3-^ uptake rates were highest in the NP and NPSi treatments, particularly during chlorophyll peaks on Days 3–4. Single-nutrient treatments showed limited changes, while uptake in the control remained low or near zero throughout the incubation. An initial increase in SiO_2_ uptake was observed across all treatments, with the highest rates occurring in the NPSi treatment in 2023. These patterns were further quantified by fitted uptake kinetics (Fig. S7). Estimated maximum uptake rates (*V*_*max*_) were 11.25 µM d^−1^ for DIN, 1.73 µM d^−1^ for PO_4_^3-^, and 28.4 µM d^−1^ for SiO_2_, with corresponding half-saturation constants (*K*_*S*_) of 12.69 µM, 0.43 µM, and 6.32 µM, respectively. Evaluation with the Willmott Skill Index showed skill values of 0.86 for DIN, 0.73 for PO_4_^3-^, and 0.67 for SiO_2_, with values closer to 1 indicating better agreement between observed and estimated uptake rates.

### Chl a dynamics in total and size-fractionated phytoplankton samples

The average Chl *a* concentration at the beginning of the experiment was 0.55 µg L^-1^ in 2022 and 0.65 µg L^-1^ in 2023. In both experiments, Chl *a* increased significantly after three days of incubation, marking the onset of a bloom phase, especially in the combined nutrient treatments. Maximum concentrations were observed on Day 4 in the NP treatment (32.5 ± 1.8 µg L^-1^ in 2022 and 14.53 ± 1.8 µg L^-1^ in 2023) and the NPSi treatment (28.35 ± 4.64 µg L^-1^), followed by a rapid decline (Fig. 4). In 2022, all treatments, including the control, exhibited significant growth by Day 3 compared to initial conditions. However, Control and P treatments subsequently declined, while the N treatment maintained constant levels from Day 3 to Day 7. In contrast, in 2023, the control, N, and P showed little change during the first half of the experiment, followed by moderate increases on Day 5, reaching maximum concentrations on Day 7 (Control: 1.3 ± 0.36 µg L^-1^; N: 2.27 ± 0.5 µg L^-1^; P: 1.6 ± 0.48 µg L^-1^).

**Figure 4.**
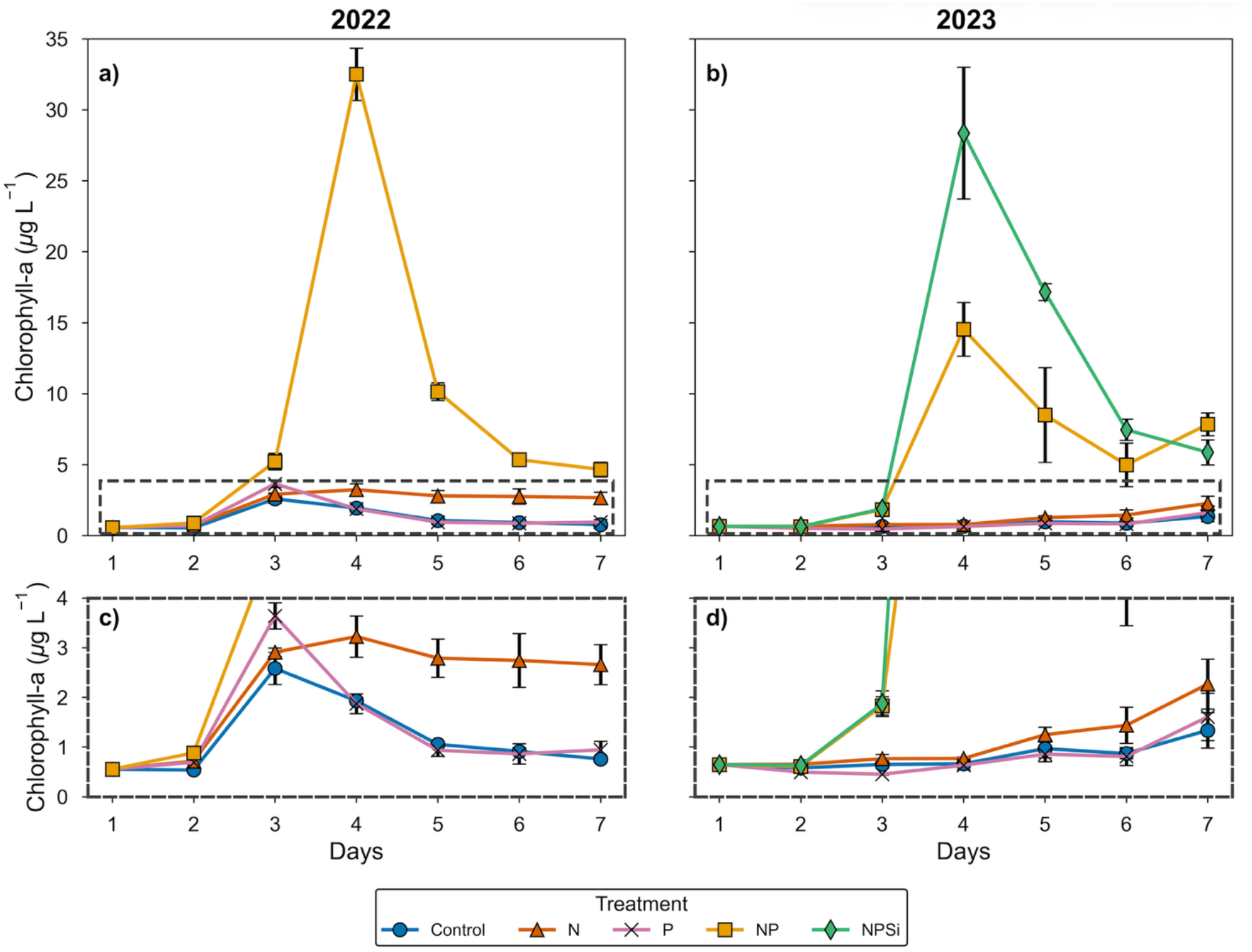
Mean Chlorophyll *a* (Chl *a*) concentrations over the 7-day incubation period under different nutrient treatments in 2022 (a, c) and 2023 (b, d). Panels (c, d) show a zoomed-in view for lower concentrations. Treatments include Control (blue), N (orange), P (red), NP (yellow), and NPS (green). Error bars represent standard error (± SE) calculated from independent replicates (n = 3)

Size-fractionated Chl *a* samples revealed significant shifts in phytoplankton community structure in response to nutrient additions and incubation time (Fig. 5). In the 2023 experiment, the initial phytoplankton community of Shiraho Reef was dominated by microphytoplankton (78.1% of total Chl *a*), followed by nanophytoplankton (15.6%) and picophytoplankton (6.3%). During the early phase of the experiment, responses were primarily driven by picophytoplankton. Tukey-adjusted comparisons showed a significant increase in their Chl *a* concentration during the first three days across all nutrient-enriched treatments, whereas no significant change was observed in the control. Following this initial rise, picophytoplankton proportions gradually declined during the bloom phase associated with combined nutrient additions. In the NP treatment, picophytoplankton increased again toward the end of the incubation, while in the NPSi treatment their contribution remained low after microphytoplankton became dominant. Nanophytoplankton showed relatively stable dynamics in the control and single nutrient treatments throughout the experiment. In contrast, during the bloom phase of the NP and NPSi treatments, nanophytoplankton proportions decreased as microphytoplankton increased and dominated Chl *a*. Microphytoplankton dominated the initial community and showed the strongest response to combined nutrient treatments. Significant increases in microphytoplankton Chl *a* were observed only in the NP and NPSi treatments, with a pronounced dominance on Day 4. After the bloom peak, microphytoplankton declined in the NP treatment but remained consistently dominant in the NPSi treatment through the end of the incubation. Total and relative contributions of pico-, nano-, and micro-sized fractions from size-fractionated Chl *a* samples are presented in Fig. 5.

**Figure 5.**
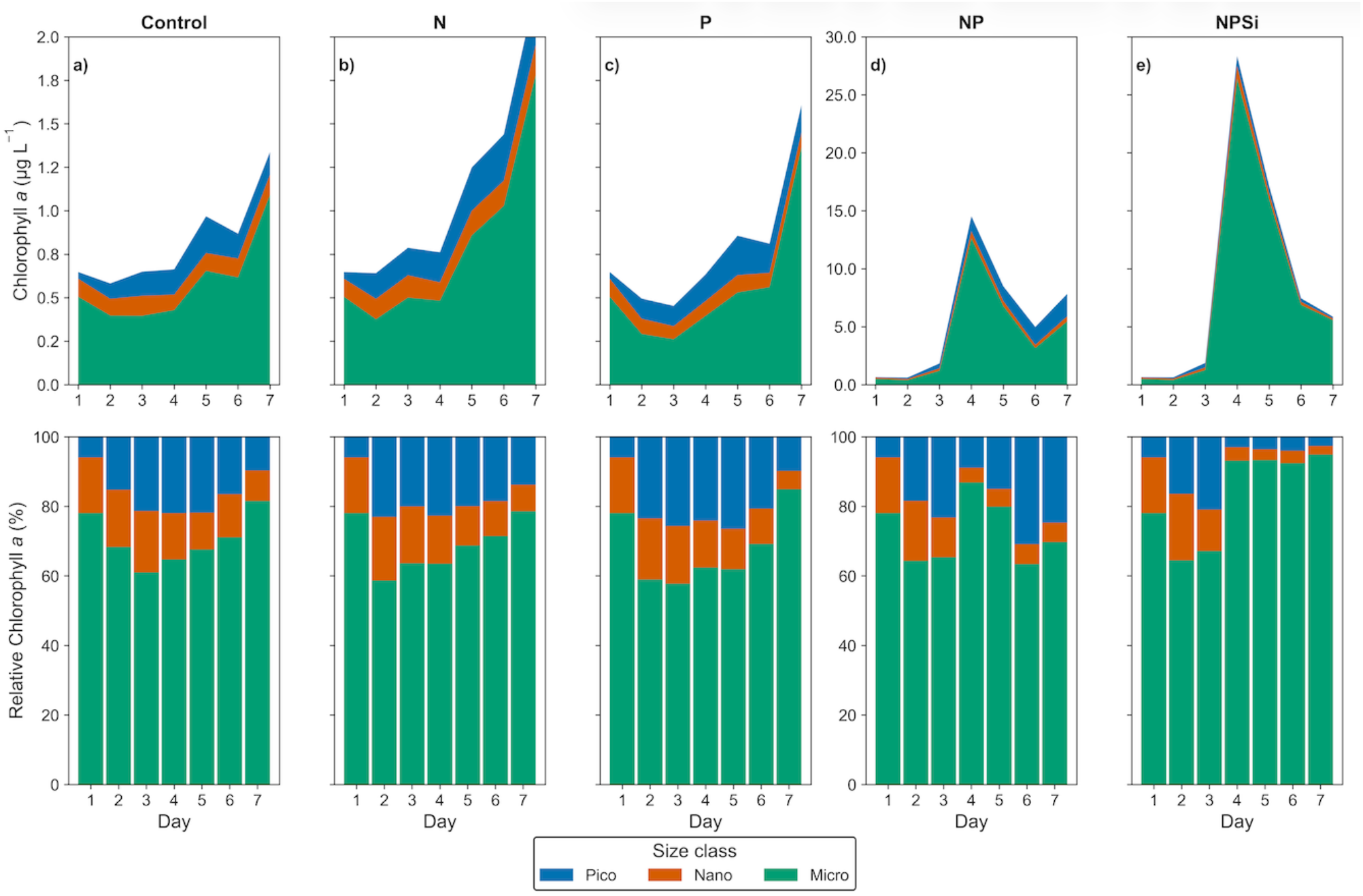
Chlorophyll *a* (Chl *a*) concentration and size-fractionated distribution over the 7-day incubation under different nutrient treatments. Top panels (a–e) show total Chl *a* concentration (µg L^−1^) for each treatment: Control (a), N (b), P (c), NP (d), and NPSi (e). Note that NP and NPSi treatments use different y-axis scales. Lower panels display relative contributions (%) of various phytoplankton size classes (picophytoplankton in blue, nanophytoplankton in orange, and microphytoplankton in green) to total Chl *a*.

### Dissolved Organic Carbon

During the first 24 h of the experiment, DOC concentrations remained stable and without significant differences between treatments in both years (Fig. 6; Table S3). After Day 3, DOC dynamics varied among treatments and years, with significant differences emerging toward the end of the incubation (Tables S13, S14). In 2022, DOC exhibited sharp fluctuations. An early peak was observed in the control treatment on Day 3, followed by higher concentrations under nutrient-enriched conditions on Day 4, particularly in the P (151.23 µM) and NP (145.03 µM) treatments. These peaks were short-lived, with concentrations declining by Day 5. Most treatments returned to near-initial concentrations by Day 6, before increasing again on Day 7. However, DOC in the NP treatment showed a secondary increase, reaching the highest overall concentration on Day 7 (166.54 µM).

**Figure 6.**
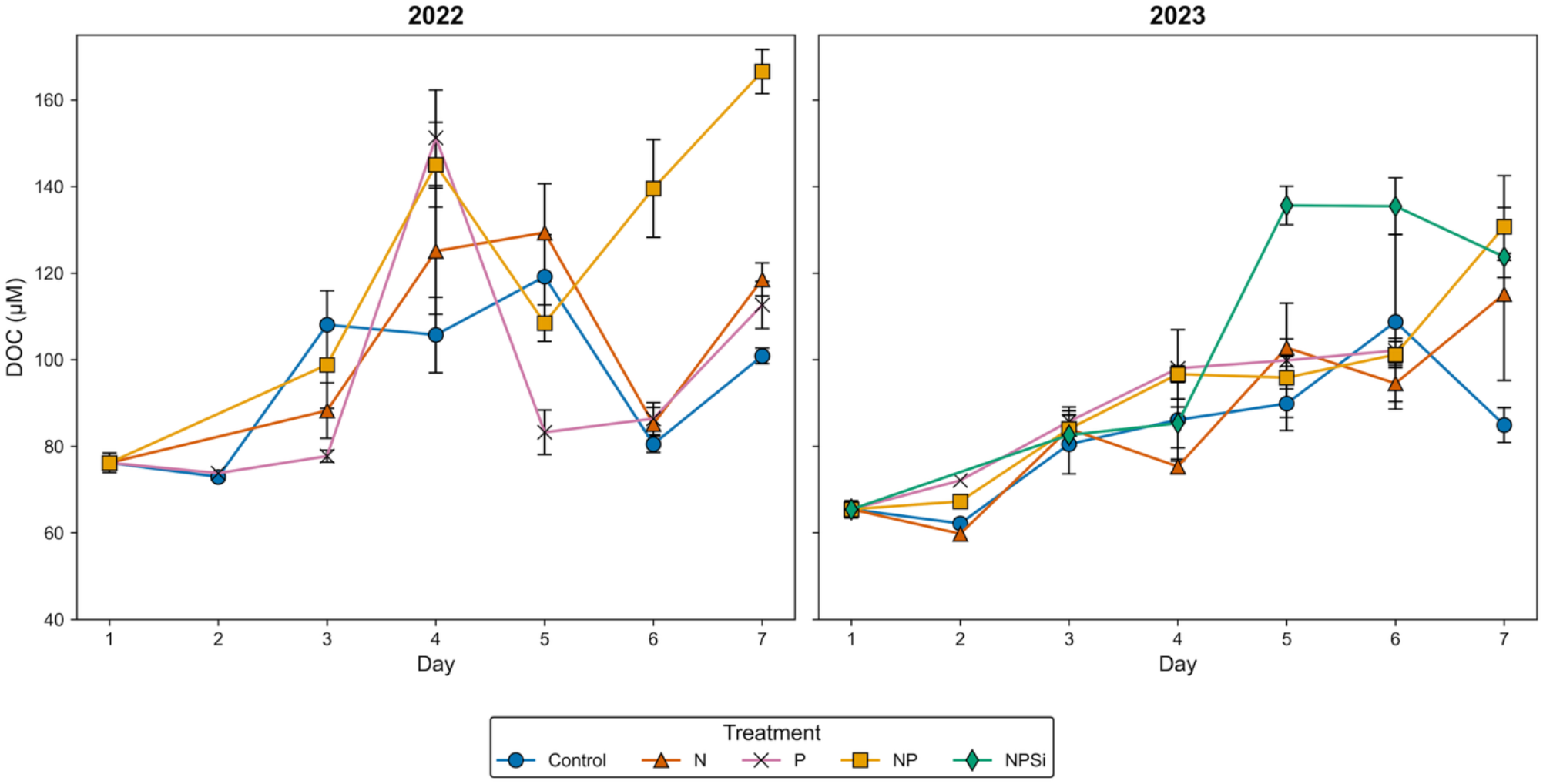
Dissolved Organic Carbon (DOC) concentrations (µM) among treatments during 7-day incubation experiments of 2022 and 2023. Colored lines indicate mean DOC concentrations for each treatment. Error bars represent standard errors of the mean (n = 3).

In 2023, DOC dynamics were more gradual and less variable. No significant differences were found between treatments until Day 5, when DOC increased sharply in the NPSi treatment. The N, P, and NP treatments showed slower increases and reached their maximum concentrations near the end of the experiment. By Day 7, DOC concentrations in NP and N exceeded those of the control, which increased gradually through Day 6 before declining on the final day to near initial concentrations.

### Carbonate system parameters

Carbonate system parameters, including bicarbonate ion (HCO_3-_), carbonate ion (CO_3_^2-^), and *p*CO_2,_ showed clear treatment-dependent trends during the 7-day incubation experiments. While control and individual nutrient additions (N and P) remained relatively constant throughout the experiment, combined nutrient treatments (NP and NPSi) led to pronounced shifts in carbonate chemistry, particularly between Days 3 and 5 (Fig. 7). In the NP and NPSi treatments, TA concentrations declined after Day 2, reaching a minimum of 2199.1 µmol kg^−1^ in the NPSi treatment by Day 5 in 2023. These changes coincided with peak Chl *a* concentrations. A similar pattern was observed in DIC, with initial values remaining stable across all treatments, followed by sharp declines in the NP and NPSi treatments after Day 3. The lowest DIC value was observed in the NPSi treatment in 2023 (1727.9 µmol kg^−1^, Day 5), followed by the NP treatment (1875.8 µmol kg^−1^).

**Fig. 7.**
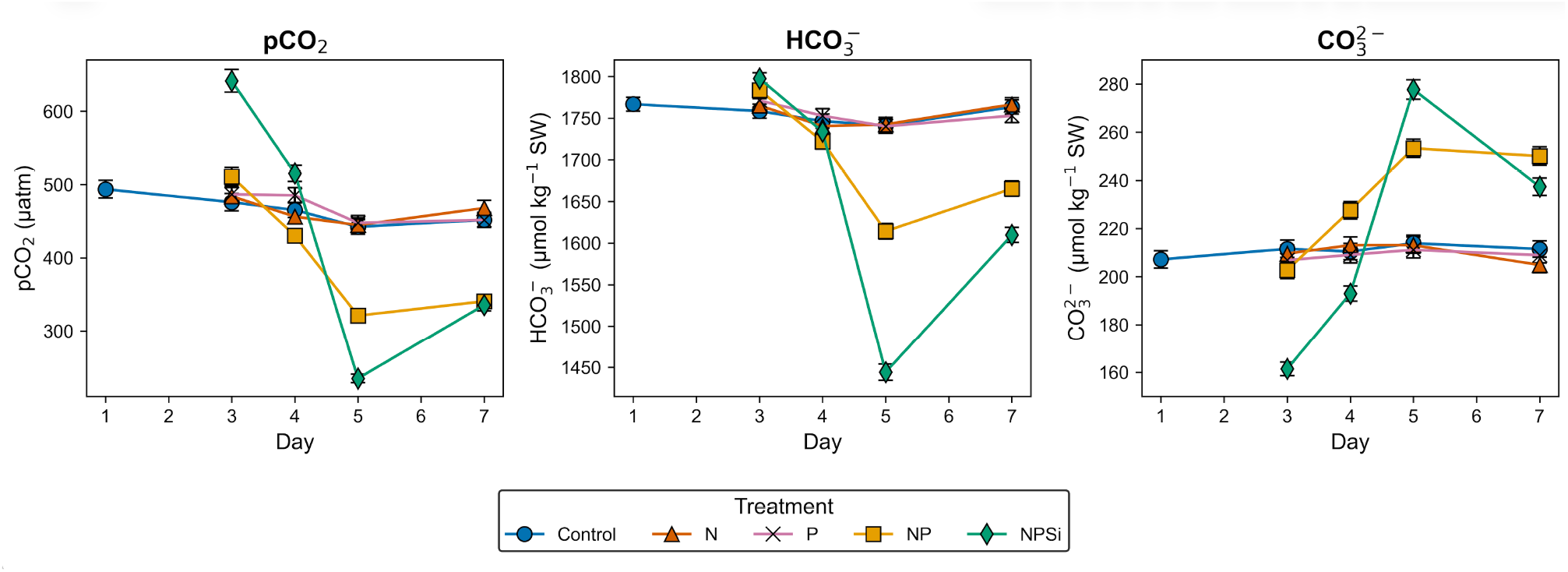
Temporal variation in carbonate system parameters during the phytoplankton incubation experiment in 2023 under different nutrient enrichment treatments. Error bars represent the root-mean-square (RMS) of propagated uncertainties derived from pH and TA measurements. Complete data and other carbonate system parameters are available in Supplementary Material (Table S4; Fig S8).

In the NP and NPSi treatments, *p*CO_2_ decreased sharply by Day 5, reaching minimum values around 233 µatm in NP (2022), 321.1 µatm in NP (2023), and 234.5 µatm in NPSi (2023). These declining trends coincide with peak Chl *a* biomass after nutrient enrichment. Toward the end of both experiments (Days 6–7), *p*CO_2_ increased again. The distribution of carbonate species in the DIC pool remained relatively constant in the control and single-nutrient treatments, with HCO_3_^−^ consistently representing ∼86–89%, CO_3_^2−^ around 10–14%, and CO_2_ <1% (Table S5). In contrast, NP and NPSi treatments exhibited shifts in carbonate speciation, with decreases in HCO_3_^−^ and corresponding increases in CO_3_^2−^. While pH_T_ values increased in combined treatments, control and individual nutrient treatments (N and P) showed no significant changes.

### Reference samples around Ishigaki Island

Chl *a* samples collected around Ishigaki Island in September 2023 (Fig. 1), revealed total Chl *a* concentrations below 0.64 µg L^-1^ in reef areas, while river-influenced areas exhibited higher values (Table S6). Size-fractionated Chl *a* further showed differences in phytoplankton size structure between river-influenced and reef environments, with microphytoplankton dominating near the river mouth and decreasing toward the reef crest (Fig. S9). Among all sampling sites, the Todoroki River mouth exhibited the highest nutrient concentrations, with DIN (10.25 µM), PO_4_^3−^ (1.01 µM), and SiO_2_ (33.85 µM). In contrast, reef crest sites, such as Shiraho Reef and Fukido Reef showed substantially lower values, with DIN ranging from 0.70 –1.56 µM and PO_4_^3−^ from 0.12–0.23 µM, reflecting low N:P ratios (∼6–8).

## Discussion

Nutrient enrichment in Shiraho Reef microcosm experiments produced rapid and consistent changes in Chl *a* concentrations, phytoplankton size structure, and carbonate system parameters. These results demonstrate strong short-term pelagic responses to episodic nutrient inputs in an otherwise oligotrophic coral reef system. In particular, combined nutrient additions resulted in elevated Chl *a* concentrations and a shift toward larger phytoplankton size classes, with microphytoplankton becoming dominant under nutrient-enriched conditions. The environmental conditions and phytoplankton responses are discussed below.

### Temperature and PFD

While light conditions remained similar in both years, temperatures recorded in 2023 were significantly higher than in 2022. It is possible that reduced precipitation and the absence of typhoons in September 2023, which cool shallow reef waters (Bernardo et al. 2017), led to higher temperatures in the microcosm experiment. High temperature and light intensity can influence biological processes, such as photoacclimation and enzymatic activity, and significantly affect growth rates and chlorophyll *a* concentrations (Geider 1987), leading to differences in community structure between experiments. In contrast to other studies that have conducted microcosm experiments under controlled conditions (Anderson *et al*., 2022; Chen *et al*., 2022), our *in-situ* experimental design maintained natural conditions, offering more realistic phytoplankton responses under natural light and temperature conditions in the field. This is particularly important in coral reefs like Shiraho, which are strongly influenced by tidal cycles and land-sea interactions (Kayanne et al. 1995; Hata et al. 2002). During our experiment in the summer of 2023, light intensity reached 2,500 µmol photons m^−2^ s^−1^ and temperatures attained 32.5 °C, which reflects the extreme conditions of reef flats and reinforces the importance of studying these systems under their natural regimes. Furthermore, these high temperatures reflect possible future scenarios in other shallow areas, which must be considered in the context of climate change.

### Chl a dynamics in total and size-fractionated phytoplankton samples

Total chlorophyll *a* concentrations showed clear patterns of phytoplankton growth in response to nutrient enrichment, particularly under combined nitrogen and phosphorus additions (NP and NPSi), in contrast to limited change with single-nutrient additions (N and P). Although chlorophyll *a* was lower in 2023, possibly due to photoacclimation or metabolic effects under higher temperatures, trends remained the same in both years. By examining size-fractionated chlorophyll *a*, significant differences in the absolute and relative contributions of pico, nano, and microphytoplankton among treatments and sites revealed ecological responses that remained hidden at the total Chl *a* level. Our results indicate that in summer, the phytoplankton community in Shiraho Reef is dominated by microphytoplankton, rather than the pico- and nanophytoplankton fractions expected in oligotrophic tropical systems. While low-nutrient environments generally select small, fast-growing cells with high surface-area-to-volume ratios that optimize nutrient uptake (Marañón et al. 2015), localized nutrient enrichment can shift the community toward larger cells, capable of luxury uptake and nutrient storage (Litchman and Klausmeier 2008). In oligotrophic coral reefs around Okinawa, pico- and nanophytoplankton are dominant contributors to chlorophyll *a* biomass (Tada et al. 2003). In contrast, our results showed a higher contribution from microphytoplankton in Shiraho Reef at the start of the experiment (∼78 %), suggesting that the reef is influenced by nutrients that promote community shifts toward larger size classes capable of nutrient storage (Yi et al. 2014; Marañón et al. 2015; Böttjer-Wilson et al. 2021; Groß et al. 2022). Particularly, microphytoplankton contributed ∼85% near the Todoroki River mouth, compared to ∼48% at the reef crest (Fig. S9). While initial nutrient concentrations and salinity did not clearly indicate influence from river and submarine groundwater inputs, dilution effects, ephemeral pulses, rapid assimilation, and unevenly distributed microenvironments can obscure their presence. Therefore, the phytoplankton size structure may represent a more sensitive indicator of nutrient limitation status in coral reefs, revealing ecological changes that may be missed when relying only on discrete measurements of total chlorophyll *a*, dissolved nutrients, or stoichiometry ratios, as suggested by recent studies (Chorus and Spijkerman 2021; Domingues et al. 2023).

### Dissolved inorganic nutrients availability, uptake, and limitation

The Todoroki River has previously been identified as a significant nutrient source for Shiraho Reef, especially nitrate (Blanco et al. 2010; Yamazaki et al. 2011). Supporting these observations, our results suggest that nitrogen limitation alone does not explain phytoplankton responses in the summer of 2022 and 2023. Although initial N:P ratios (∼11.6 in both 2022 and 2023) fall below the classical Redfield ratio (16:1; Redfield, 1934) and align with the standard paradigm of nitrogen limitation in coastal and coral reef environments (Vitousek et al. 1997; Downing et al. 1999), experimental responses to nutrient additions revealed more complex dynamics. The modest reduction in DIN under N-only addition and minimal phytoplankton response to P alone contrasts sharply with the strong DIN uptake and biomass increase in the NP and NPSi treatments. These patterns indicate conditional co-limitation, in which the growth response to one nutrient depends on the availability of another (Elser et al. 2007; Harpole et al. 2011). Consistent with this, uptake kinetics (*Vmax* and *K*) indicate that nutrient concentrations in summer were below half-saturation thresholds, limiting efficient uptake unless both nutrients were supplied simultaneously. In contrast, silicate exhibited a high *Vmax* and an intermediate *K*, indicating a high SiO_2_ uptake capacity when N and P were supplied simultaneously. These patterns support a framework of conditional co-limitation, likely influenced by nitrogen enrichment from terrestrial sources. Therefore, runoff from the Todoroki watershed in summer likely increases DIN concentrations, contributing to N and P co-limitation in the reef system.

Submarine groundwater discharge (SGD) is another key nutrient source of nitrate and silicate to Shiraho Reef (Umezawa et al. 2002), providing a constant, but spatially heterogeneous nutrient source that may support phytoplankton even under low riverine flow. On the other hand, phosphorus is transported via surface runoff associated with erosion events (Ikeda et al. 2009). Water residence times in Shiraho Reef (3-8 h; Yamano et al. 2014) are short, yet sufficient to allow nutrient accumulation, especially during low tides when exchange with offshore waters is reduced, and nutrients accumulate in the shallow water column. Episodic freshwater inputs during the rainy and typhoon seasons can further modify nutrient availability and drive shifts in the phytoplankton community structure within the reef (Blanco et al. 2008; Tedetti et al. 2020).

In this context, size-fractionated Chl *a* is an effective indicator of nutrient regimes across reef environments. As shown by the comparative analysis of various sites around Ishigaki Island, size-fractionated Chl *a* revealed the dominance of smaller phytoplankton fractions at reef crests. In contrast, sites under direct riverine influence, e.g., Todoroki and Nagura, exhibited higher proportions of larger cells, indicating increased nutrient availability. Therefore, phytoplankton size structure provides a more integrative proxy for ecosystem status than nutrient concentrations alone, which are often diluted or missed by discrete sampling due to temporal or spatial heterogeneity.

### Phytoplankton-driven carbon dynamics

Dissolved organic and inorganic carbon dynamics observed in these microcosm experiments demonstrate a significant, yet underappreciated role of pelagic organisms in coral reef biogeochemistry (Shakya and Allgeier 2023). In both years, nutrient-enriched conditions directly enhanced phytoplankton growth, leading to higher production and accumulation of DOC, a decrease in *p*CO2, and shifts in carbonate speciation. These changes reflect a tightly coupled system in which planktonic processes shape carbon fluxes in the water column on short timescales.

In 2022, DOC concentrations showed sharp, short-term fluctuations in the NP and P treatments, peaking around Day 4, in parallel with chlorophyll *a*. This suggests a pulse of dissolved organic matter (DOM), likely exuded by phytoplankton during rapid growth, microzooplankton grazing and excretion, and subsequently consumed by heterotrophic bacteria (Marañón et al. 2015). Rapid turnover of DOC is consistent with efficient microbial loop dynamics observed in oligotrophic reef environments (Azam and Malfatti 2007). In contrast, the 2023 experiment showed a more gradual and sustained increase in DOC across all treatments, possibly reflecting differences in microbial degradation capacity, species composition, or temperature-dependent responses, such as phytoplankton exudation of DOM, and bacterial enzymatic activity (Pomeroy and Wiebe 2001). In the 2023 experiment, the NPSi treatment relieved silicate limitation, leading to higher DOC levels, consistent with the activity of diatoms and silico-flagellates, which release significant amounts of extracellular organic matter, including colloidal and semi-labile DOM fractions (Behrendt et al. 2024).

Significant changes in carbonate system parameters indicated elevated photosynthetic activity, especially in the NP and NPSi treatments. These changes were characterized by sharp declines in DIC, elevated pH, and a shift in carbonate speciation from HCO_3_^−^ to CO_3_^2−^ (Zeebe and Wolf-Gladrow 2001; Riebesell et al. 2007). TA remained stable in most treatments, but declined in NP and NPSi following bloom peaks. This decrease, together with increases in DIC and *p*CO_2_, suggests remineralization of organic matter driven by microbial respiration processes (Azam et al. 1983). These observations reflect the cyclic nature of carbon uptake and release, mediated by phytoplankton growth and microbial degradation in the water column, especially after phytoplankton bloom collapse. In the NPSi treatment, the most substantial shifts in TA and DIC were rapidly reversed as the bloom declined, suggesting that while photosynthetic activity increases CO_3_^2−^ and raises pH during active growth, microbial respiration of bloom-derived organic matter releases CO_2_ and drives rebound acidification (Duarte et al. 2013). These dynamics were captured despite constraints of our experimental design, which used once-daily sampling, and although we did not resolve diel variability as in Hata et al. (2002), we observed net changes that indicate cumulative biological effects following nutrient enrichment. Incorporating these pelagic and microbial processes into biogeochemical models will be critical to accurately predict coral reef metabolic responses to nutrient enrichment and climate variability.

Despite the traditional emphasis on benthic macro-organisms in reef carbon budgets, our findings highlight the central role of biological processes in the water column regulating reef-scale CO_2_ fluxes. Nutrient enrichment stimulated phytoplankton growth, enhancing short-term CO_2_ uptake and temporarily turning Shiraho Reef water into a CO_2_ sink, as indicated by significant drawdown of *p*CO_2_. Concurrent increases of chlorophyll *a* and DOC, together with previous evidence of offshore POC export (Hata et al., 2002), suggest that Shiraho Reef may function as an episodic carbon sink under certain conditions, particularly when larger phytoplankton dominate. Yet, post-bloom respiration and DOC remineralization by heterotrophic bacteria can contribute to elevated *p*CO_2_ and localized acidification, influencing trophic transfer and carbon sequestration potential (Haas et al. 2016; Nakajima et al. 2017; Nelson et al. 2023), contributing to the ongoing debate over whether coral reefs act as net sources or sinks of CO_2_ (Kayanne 2025). Our study demonstrates that phytoplankton communities may significantly influence carbon cycling and carbonate chemistry under nutrient-enriched conditions. An integrative understanding of these dynamics is essential to predict coral reef resilience in the face of climate change and increasing anthropogenic pressures.

This study has several limitations common to short-term microcosm experiments. Daily subsampling reduced experimental volume, and the exclusion of mesozooplankton may have diminished top-down control and other trophic interactions (Nogueira et al. 2014). In addition, Na_2_SiF_6_, used as a silicate source in the NPSi treatment, induced an immediate, non-biological decrease in pH, likely due to hydrolysis of hexafluorosilicate (SiF_6_^2−^) in solution (Finney et al. 2006). Such chemical perturbations directly alter carbonate system parameters and may influence phytoplankton responses independently of nutrient availability. Although Na_2_SiF_6_ has been previously used in nutrient enrichment experiments (Domingues et al. 2011), the associated pH changes are not commonly documented. Future experiments should use sodium metasilicate (Na_2_SiO_3_) as a more inert alternative (Ortiz et al. 2024; Ferderer et al. 2024). Chlorophyll *a* was used as the primary proxy for phytoplankton biomass in this study, despite known variability in pigment content across taxa and environmental conditions (Kruskopf and Flynn 2006; Ara et al. 2019; Anugerahanti et al. 2021). Nevertheless, size-fractionated chlorophyll *a* remains a simple yet effective method to characterize the size structure of phytoplankton communities and offers valuable insights into nutrient availability and ecosystem dynamics in coral reef areas. Future studies integrating size-fractionated Chl *a* with taxonomic, trait-based, and optical approaches are needed to resolve species-level shifts and improve our understanding of ecosystem responses to changing nutrient regimes.

## Conclusion

This study revealed how episodic nutrient enrichment can significantly alter biological and biogeochemical processes in water columns of coral reef ecosystems. In oligotrophic systems like Shiraho Reef, sudden increases in nutrient availability can shift phytoplankton community size structure, favoring larger cells and triggering rapid changes in nutrient cycling and carbon dynamics at the base of the food web. Shiraho Reef remained primarily nitrogen-limited during the summers of 2022 and 2023, but the biological response to nitrogen addition was strongly amplified when phosphorus was simultaneously supplied and further intensified with silicate availability, indicating dynamic co-limitation in the area influenced by terrestrial nutrient runoff. Moreover, our results show that size-fractionated chlorophyll *a* provides ecologically meaningful information about nutrient status and can serve as a sensitive bioindicator of recent nutrient pulses in reef systems where dissolved nutrients are often undetectable due to rapid uptake or dilution effects. The summer phytoplankton community in Shiraho Reef was dominated by microphytoplankton, rather than the expected pico- and nanophytoplankton dominance, typical in tropical oligotrophic systems. This indicates favorable growth conditions for larger cells driven by riverine and groundwater nutrient inputs to the reef, despite low measured dissolved nutrient levels. As climate change intensifies rainfall and storm events, these nutrient inputs will likely become a more important driver of phytoplankton communities and biogeochemical cycling. Further work should focus on quantifying nutrient runoff and its effects on marine food webs, reef resilience, and coupled interactions between planktonic and benthic components in coral reefs. Understanding nutrient co-limitation dynamics will be essential for developing effective management strategies to safeguard biodiversity, ecosystem services, and reef health in a changing world.

## Statements & Declarations

## Acknowledgements

The authors thank Dr. Satoshi Mitarai for his invaluable advice and full support during the preparation of this paper, and the technical editor, Steven D. Aird, for his careful editorial assistance.

## Compliance with ethical standards

The authors declare that they have no conflict of interest. Approval to install the experimental setup at Shiraho Beach (Iriomote-Ishigaki National Park) was granted by the Ministry of the Environment of Japan.

## Data availability

All datasets necessary to replicate analyses are available in a GitHub repository https://github.com/georgesuac/Microcosms_Ishigaki

## Author Contributions

All authors contributed to the study conception and design. JS carried out the data collection and laboratory work. Data analysis and interpretation were performed by JS and TN. JS prepared the first draft of the manuscript. All authors reviewed and approved the final manuscript.

