## Supplementary-material for "Phytoplankton size structure and biogeochemical responses to nutrient enrichment in an oligotrophic coral reef"

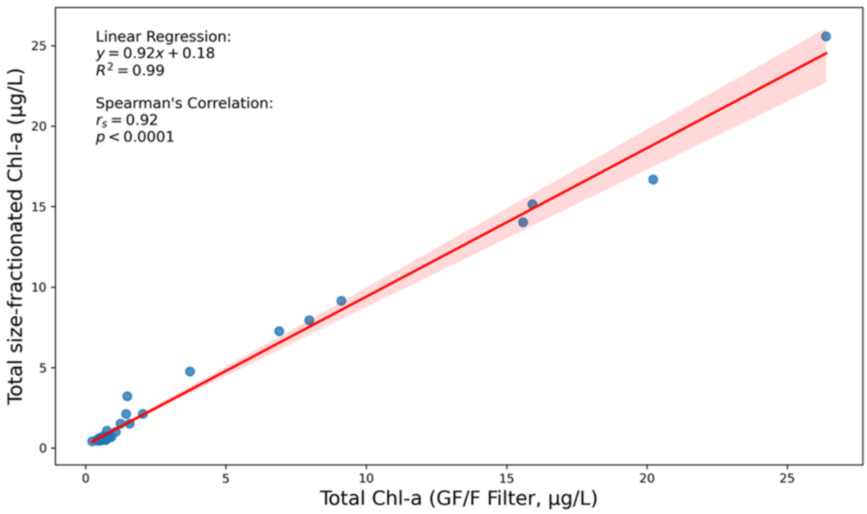


**Figure S1.** Linear correlation of total chlorophyll *a* concentrations among size-fractionated samples and GF/F filters in the experiment of 2023 (Spearman's r = 0.92, P = 2.48e$^{-17}$, N = 40). No significant differences were found between the total Chl-a concentration (GF/F filters) and the total concentration of size-fractionated samples. This demonstrates the good retention capacity of both methods. The red shaded area represents the 95% confidence interval.

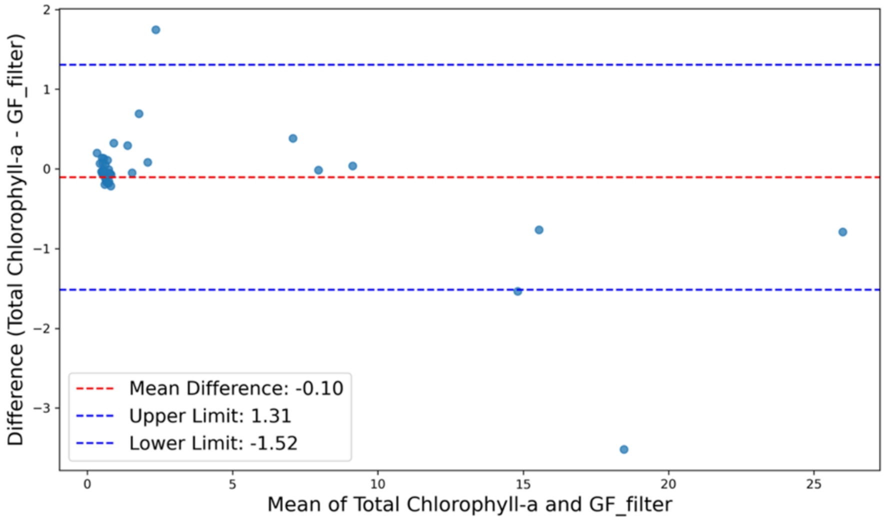


**Figure S2.** Bland-Altman Plot: Total Chlorophyll-a from size-fractionated samples vs GF filters (Wilcoxon Test Statistic, T = 320;  P = 0.4727). The 95% limits of agreement range from -1.52 to 1.31 µg/L. Most data points lie within these limits, showing that the two methods agree well overall. However, at higher Chl-a concentrations, a few points deviate slightly, indicating potential systematic differences or method limitations under high biomass conditions.

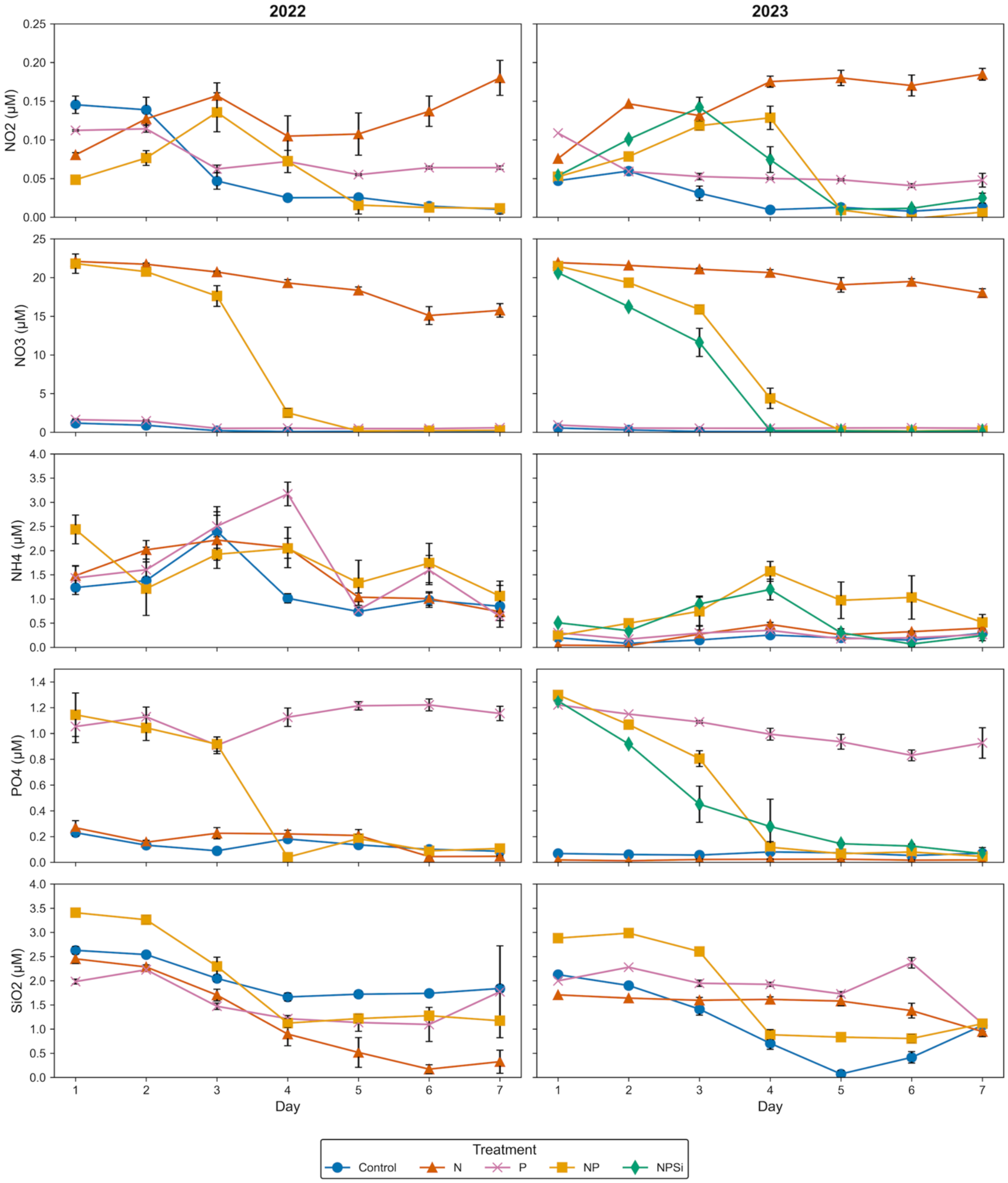


**Figure S3.** Individual dissolved inorganic nutrients measured throughout the 7-day incubation period.

a)

b)

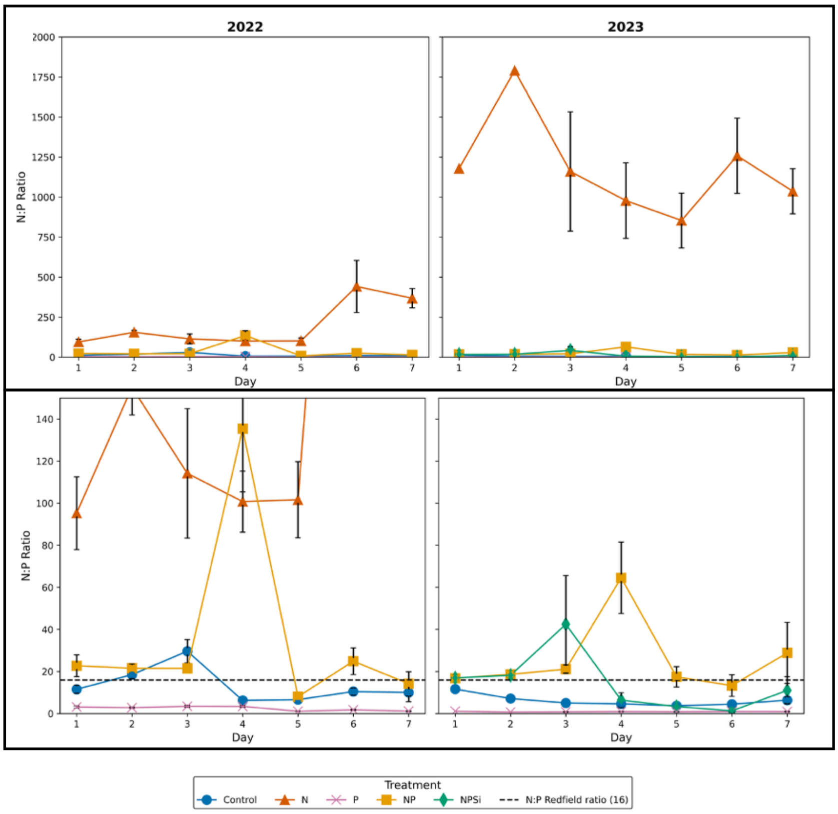


**Figure S4.** N:P ratios throughout the 7-day incubation period. The dashed line represents the Redfield Ratio. Panels in b) are close-ups of panels in a). A high N:P ratio in enriched treatments, as observed in N, NP, and NPSi treatments, may indicate that P becomes the limiting factor over time.

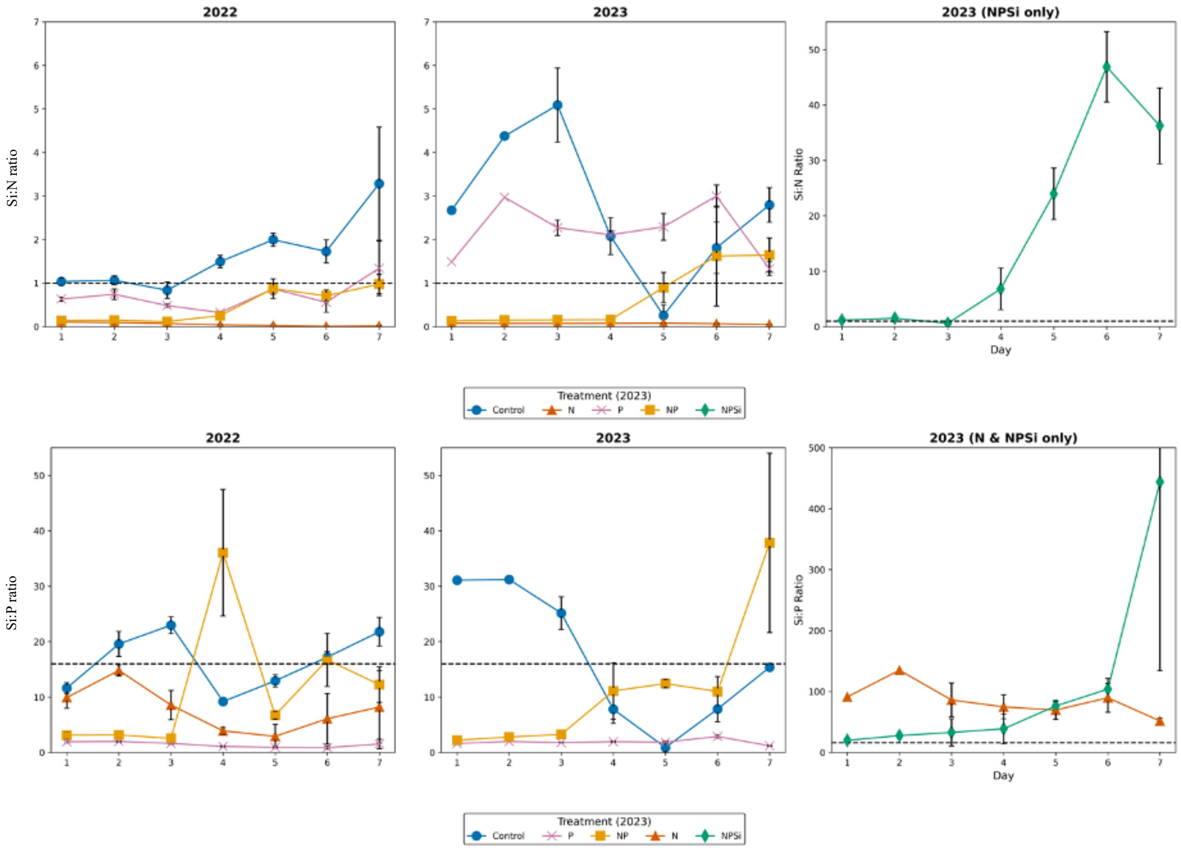


**Figure S5.** Temporal changes in the Si:N and Si:P ratios during the incubation period.

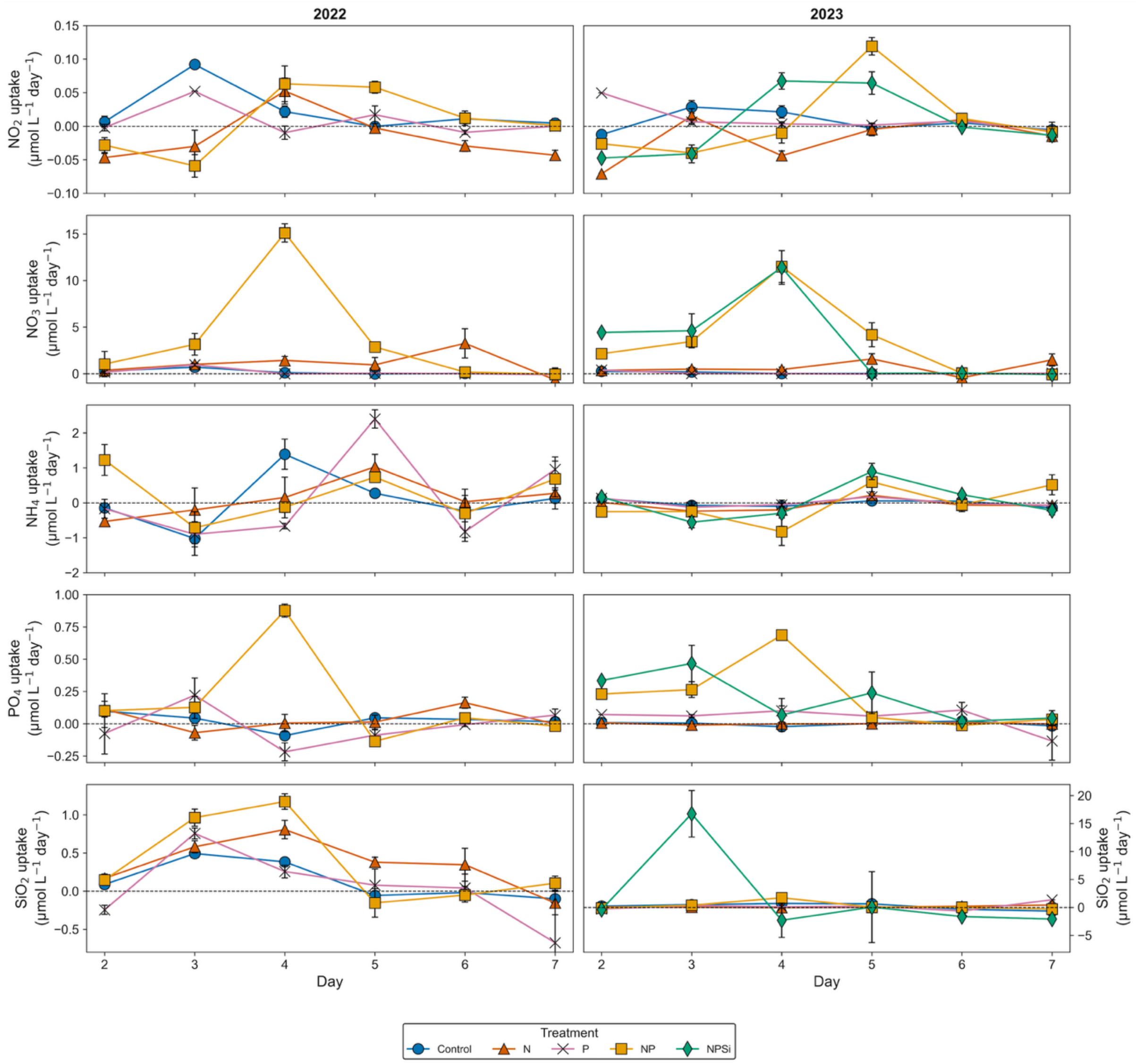


**Figure S6.** Daily nutrient uptake rates for each treatment during the experiments in 2022 and 2023, calculated from finite differences per day. Points represent mean uptake rates, and error bars represent standard error (n=3). Note that $SiO_2$ uptake in 2023 has a different scale than in 2022. The horizontal dashed line represents zero uptake.

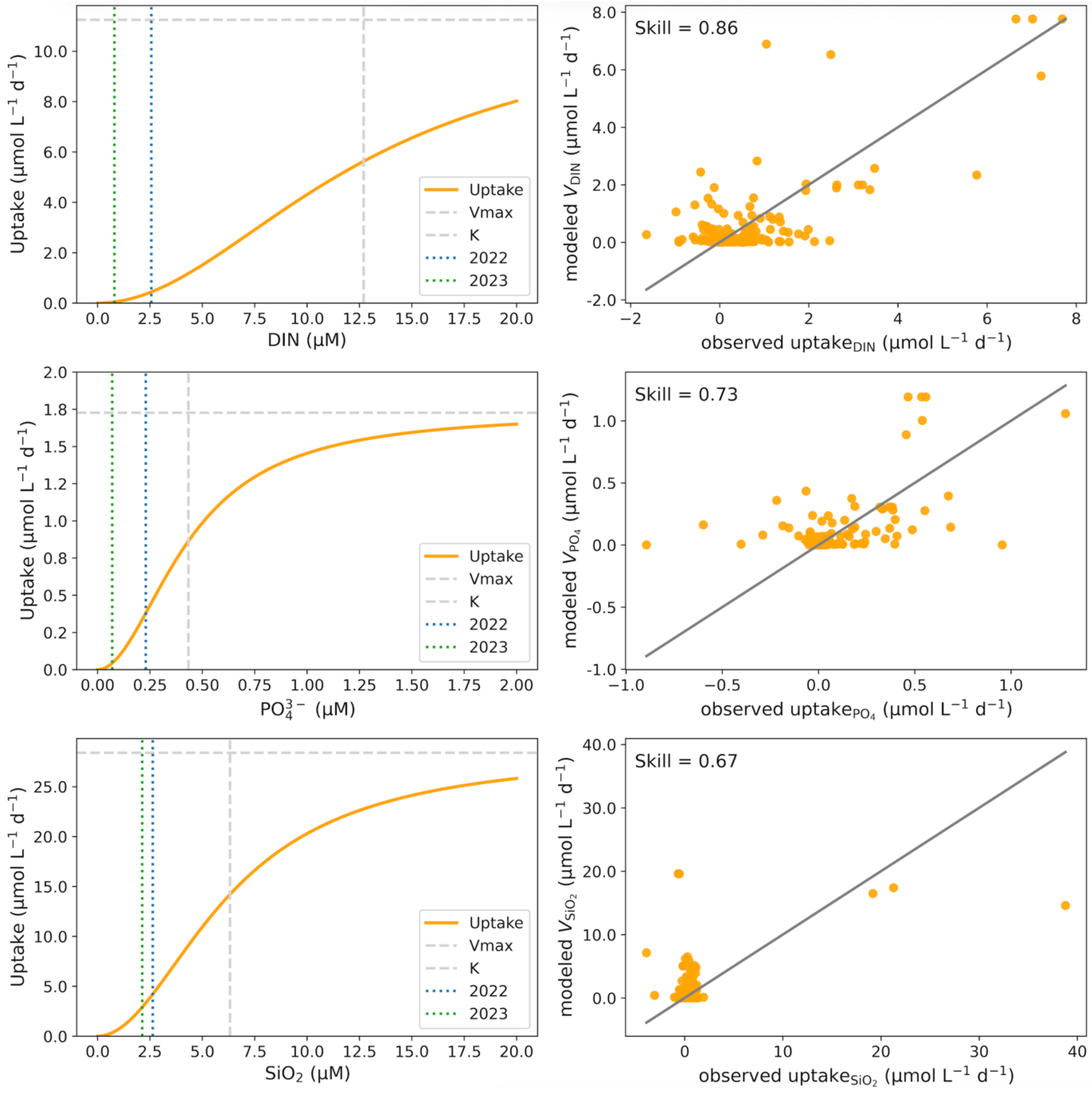


**Figure S7.** Modeled nutrient uptake kinetics for Shiraho Reef phytoplankton communities. Uptake rates were estimated using Hill-type functions with co-limitation among dissolved inorganic nitrogen (DIN), phosphate ($PO_4^{3-}$), and silicate ($SiO_2$). Panels on the left show fitted uptake curves (solid orange lines). Horizontal dashed gray lines indicate maximum uptake rates (*Vmax*), vertical dashed gray lines indicate half-saturation constants (*K*). Vertical dotted blue and green lines mark measured ambient nutrient concentrations in summer 2022 (DIN = 2.56 μM; $PO_4^{3-}$ = 0.23 μM; $SiO_2$ = 2.63 μM) and 2023 (DIN = 0.8 μM; $PO_4^{3-}$ = 0.07 μM; $SiO_2$ = 2.13 μM), respectively. Panels on the right show comparisons between observed and modeled uptake rates after optimization using the Willmott Skill Index. Skill values are reported in each panel, and solid lines represent the 1:1 relationship.

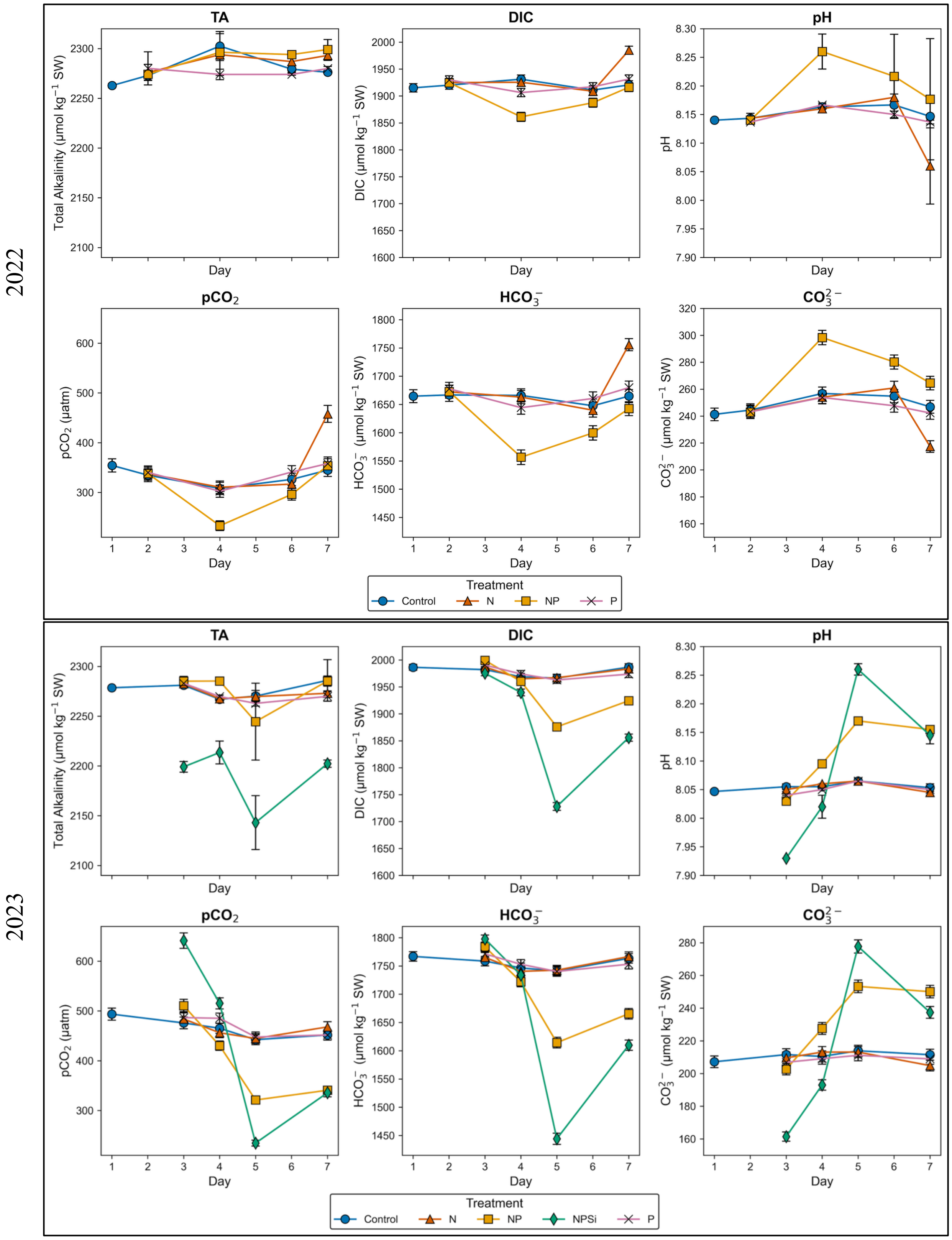


**Figure S8.** Carbonate System Parameters in the 2022 and 2023 experiments. Dissolved inorganic carbon (DIC), partial pressure of carbon dioxide ($pCO_2$), bicarbonate ion ($HCO_3^-$), carbonate ion ($CO_3^{2-}$), and dissolved carbon dioxide ($CO_2$) were based on measured $pH_T$ (total scale) and total alkalinity (TA). For TA and pH, error bars represent the standard deviation of replicate measurements (n=2-3). For all other parameters, error bars represent the root-mean-square (RMS) of propagated errors calculated with CO2SYS version 3.0 (Lewis et al. 1998). The y-axis scale is the same within parameters for better comparability between years.

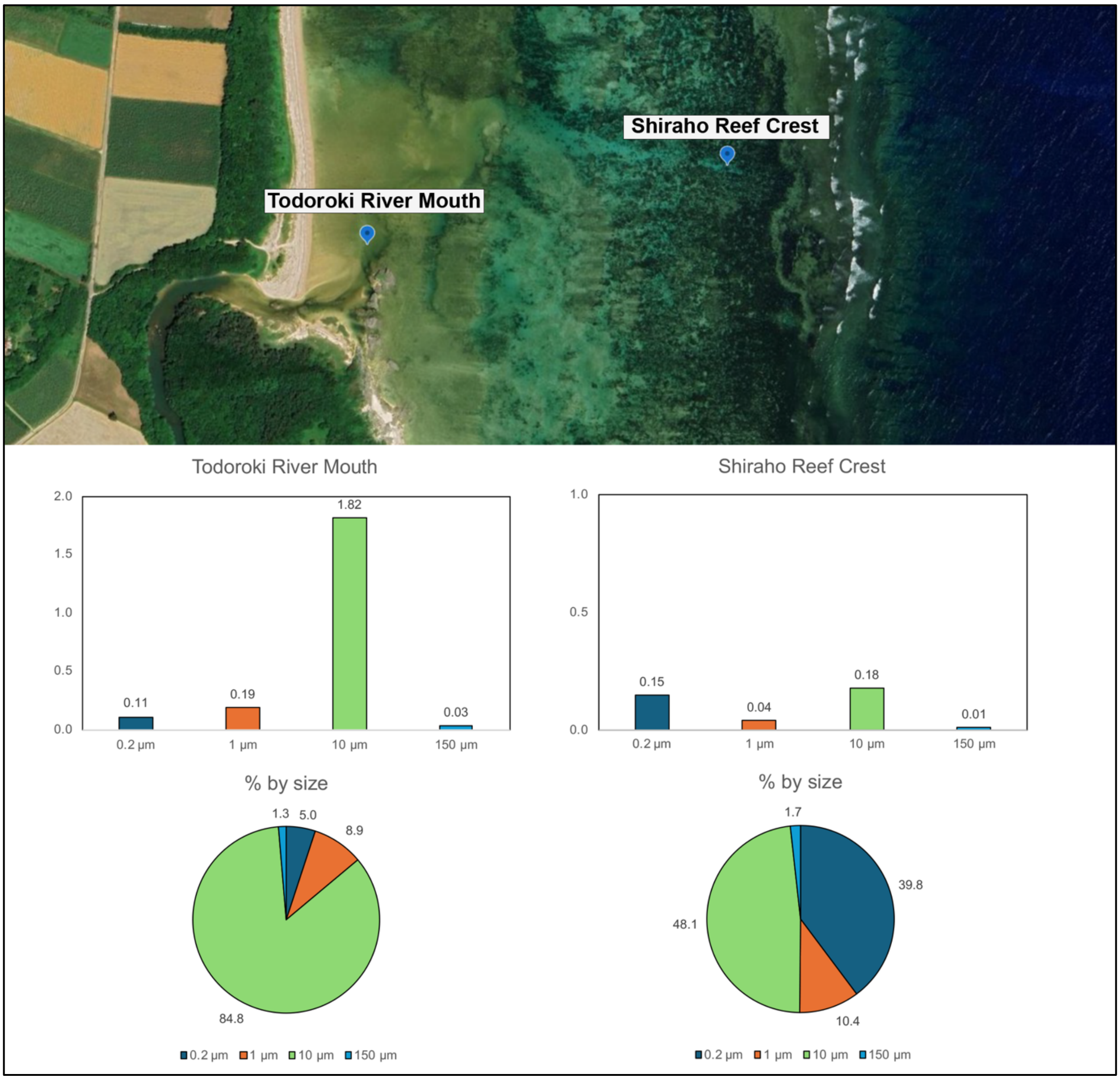


**Figure S9.** Spatial comparison of size-fractionated Chlorophyll *a* (Chl *a*) concentrations between Todoroki River Mouth and Shiraho Reef Crest, using the same size-fractionation method applied in the microcosm experiments (0.2-1, 1–10, and 10–150 µm size fractions). Microphytoplankton (10-150 µm) strongly dominated near the river mouth (84.8%), whereas their contribution decreased at the reef crest (48.1%). This gradient shows differences in phytoplankton size structure associated with local nutrient conditions. (Satellite image: Google Earth).

**Table S1.** Michaelis-Menten-type co-limitation functions used to estimate uptake rates of DIN, $PO_4^{3-}$, and SiO2 in nutrient enrichment experiments at Shiraho Reef, including Willmott Skill Index used to evaluate model performance.

| Variable/Index | Formula |
|---|---|
| DIN uptake | $V_{DIN} = V_{DIN\_max} \min\left(\frac{DIN^2}{DIN^2 + K_{DIN}{}^2}, \frac{PO_4{}^2}{PO_4{}^2 + K_{PO_4}{}^2}, \frac{SiO_2{}^2}{SiO_2{}^2 + K_{SiO_2}{}^2}\right)$ |
| DIP uptake ( | $V_{PO_4} = V_{PO_4\_max} \min\left(\frac{DIN^2}{DIN^2 + K_{DIN}{}^2}, \frac{PO_4{}^2}{PO_4{}^2 + K_{PO_4}{}^2}, \frac{SiO_2{}^2}{SiO_2{}^2 + K_{SiO_2}{}^2}\right)$ |
| DSi uptake | $V_{SiO_2} = V_{SiO_2\_max} \min\left(\frac{DIN^2}{DIN^2 + K_{DIN}{}^2}, \frac{PO_4{}^2}{PO_4{}^2 + K_{PO_4}{}^2}, \frac{SiO_2{}^2}{SiO_2{}^2 + K_{SiO_2}{}^2}\right)$ |
| Willmott Skill Index (0–1);<br>1 = perfect agreement,<br>0 = no agreement | $\text{Skill} = 1 - \frac{\sum \lvert X_{model} - X_{obs} \rvert^2}{\sum \left( \lvert X_{model} - \overline{X_{obs}} \rvert + \lvert X_{obs} - \overline{X_{obs}} \rvert \right)^2}$ |

**Table S2.** Dissolved Inorganic Nutrient concentrations measured during the 7-day incubation experiments in 2022 and 2023. Values represent the mean ± standard deviation of replicate bottles (n = 2–3) for each treatment and day.

| Nutrient | | Treatment | Day 1 | Day 2 | Day 3 | Day 4 | Day 5 | Day 6 | Day 7 |
|---|---|---|---|---|---|---|---|---|---|
| $NH_4^+$ | 2022 | Control | 1.24 ± 0.25 | 1.38 ± 0.12 | 2.43 ± 0.87 | 1.01 ± 0.17 | 0.74 ± 0.08 | 0.98 ± 0.26 | 0.85 ± 0.75 |
| | | N | 1.48 ± 0.36 | 2.02 ± 0.33 | 2.22 ± 1.01 | 2.07 ± 0.73 | 1.04 ± 0.15 | 1.01 ± 0.24 | 0.73 ± 0.18 |
| | | P | 1.43 ± 0.42 | 1.6 ± 0.8 | 2.51 ± 0.39 | 3.17 ± 0.42 | 0.77 ± 0.09 | 1.6 ± 0.52 | 0.64 ± 0.15 |
| | | NP | 2.44 ± 0.52 | 1.21 ± 0.96 | 1.92 ± 0.22 | 2.05 ± 0.36 | 1.33 ± 0.66 | 1.74 ± 0.7 | 1.06 ± 0.53 |
| | 2023 | Control | 0.2 | 0.08 | 0.15 ± 0.03 | 0.25 ± 0.05 | 0.2 ± 0.06 | 0.07 ± 0.02 | 0.29 ± 0.03 |
| | | N | 0.04 | 0.03 | 0.27 ± 0.29 | 0.47 ± 0.08 | 0.26 ± 0.02 | 0.32 ± 0.03 | 0.4 ± 0.03 |
| | | P | 0.3 | 0.17 | 0.29 ± 0.09 | 0.35 ± 0.07 | 0.18 ± 0.08 | 0.2 ± 0.005 | 0.27 ± 0.04 |
| | | NP | 0.25 | 0.5 | 0.74 ± 0.51 | 1.57 ± 0.36 | 0.97 ± 0.65 | 1.03 ± 0.78 | 0.52 ± 0.28 |
| | | NPSi | 0.51 | 0.34 | 0.9 ± 0.28 | 1.2 ± 0.37 | 0.3 ± 0.14 | 0.07 ± 0.08 | 0.24 ± 0.15 |
| $NO_2^-$ | 2022 | Control | 0.15 ± 0.02 | 0.14 ± 0.03 | 0.05 ± 0.02 | 0.03 ± 0.005 | 0.03 ± 0.005 | 0.01 ± 0.01 | 0.01 ± 0.01 |
| | | N | 0.08 ± 0.01 | 0.13 ± 0.01 | 0.16 ± 0.03 | 0.1 ± 0.05 | 0.11 ± 0.05 | 0.14 ± 0.03 | 0.18 ± 0.04 |
| | | P | 0.11 ± 0.002 | 0.11 ± 0.01 | 0.06 ± 0.01 | 0.07 ± 0.03 | 0.05 ± 0.002 | 0.06 ± 0.003 | 0.06 ± 0.004 |
| | | NP | 0.05 ± 0.01 | 0.08 ± 0.02 | 0.14 ± 0.04 | 0.07 ± 0.003 | 0.02 ± 0.02 | 0.01 ± 0.001 | 0.01 ± 0.003 |
| | 2023 | Control | 0.05 | 0.06 | 0.03 ± 0.02 | 0.01 ± 0.002 | 0.01 ± 0.004 | 0.01 ± 0.002 | 0.01 ± 0.01 |
| | | N | 0.08 | 0.15 | 0.13 ± 0.02 | 0.18 ± 0.01 | 0.18 ± 0.02 | 0.17 ± 0.02 | 0.18 ± 0.01 |
| | | P | 0.11 | 0.06 | 0.05 ± 0.01 | 0.05 ± 0.002 | 0.05 ± 0.002 | 0.04 ± 0.004 | 0.05 ± 0.01 |
| | | NP | 0.05 | 0.08 | 0.12 ± 0.01 | 0.13 ± 0.03 | 0.01 ± 0.01 | 0 ± 0.001 | 0.01 ± 0.001 |
| | | NPSi | 0.05 | 0.1 | 0.14 ± 0.02 | 0.07 ± 0.03 | 0.01 ± 0.001 | 0.01 ± 0.003 | 0.02 ± 0.01 |
| $NO_3^-$ | 2022 | Control | 1.18 ± 0.06 | 0.9 ± 0.31 | 0.2 ± 0.08 | 0.09 ± 0.02 | 0.1 ± 0.02 | 0.07 ± 0.05 | 0.07 ± 0.1 |
| | | N | 22.09 ± 0.44 | 21.72 ± 0.27 | 20.72 ± 0.2 | 19.3 ± 0.72 | 18.35 ± 0.76 | 15.09 ± 2 | 15.76 ± 1.52 |
| | | P | 1.63 ± 0.16 | 1.46 ± 0.23 | 0.51 ± 0.02 | 0.53 ± 0.07 | 0.47 ± 0.03 | 0.47 ± 0.02 | 0.58 ± 0.15 |
| | | NP | 21.8 ± 2.15 | 20.77 ± 0.33 | 17.62 ± 2.32 | 2.52 ± 0.9 | 0.17 ± 0.002 | 0.21 ± 0.05 | 0.26 ± 0.02 |
| | 2023 | Control | 0.55 | 0.29 | 0.1 ± 0.01 | 0.08 ± 0.02 | 0.06 ± 0.02 | 0.1 ± 0.01 | 0.09 ± 0.04 |
| | | N | 21.93 | 21.57 | 21.08 ± 0.24 | 20.64 ± 0.67 | 19.06 ± 1.64 | 19.49 ± 0.63 | 18.01 ± 0.92 |
| | | P | 0.93 | 0.54 | 0.52 ± 0.04 | 0.52 ± 0.02 | 0.55 ± 0.09 | 0.56 ± 0.11 | 0.53 ± 0.01 |
| | | NP | 21.48 | 19.34 | 15.88 ± 0.82 | 4.39 ± 2.28 | 0.21 ± 0.05 | 0.16 ± 0.12 | 0.23 ± 0.01 |
| | | NPSi | 20.64 | 16.22 | 11.61 ± 3.17 | 0.21 ± 0.02 | 0.18 ± 0.06 | 0.1 ± 0.09 | 0.17 ± 0.004 |
| $PO_4^{3-}$ | 2022 | Control | 0.23 ± 0.05 | 0.13 ± 0.03 | 0.09 ± 0.01 | 0.18 ± 0.02 | 0.14 ± 0.02 | 0.1 ± 0.01 | 0.09 ± 0.01 |
| | | N | 0.27 ± 0.1 | 0.16 ± 0.02 | 0.23 ± 0.08 | 0.22 ± 0.05 | 0.21 ± 0.08 | 0.04 ± 0.02 | 0.05 ± 0.01 |
| | | P | 1.05 ± 0.22 | 1.13 ± 0.13 | 0.91 ± 0.11 | 1.13 ± 0.12 | 1.21 ± 0.06 | 1.22 ± 0.08 | 1.16 ± 0.1 |
| | | NP | 1.14 ± 0.29 | 1.04 ± 0.17 | 0.92 ± 0.1 | 0.04 ± 0.03 | 0.18 ± 0.05 | 0.09 ± 0.04 | 0.11 ± 0.04 |
| | 2023 | Control | 0.07 | 0.06 | 0.06 ± 0 | 0.24 ± 0.02 | 0.07 ± 0.01 | 0.13 ± 0 | 0.07 ± 0.02 |
| | | N | 0.02 | 0.01 | 0.02 ± 0.01 | 0.02 ± 0.01 | 0.02 ± 0.01 | 0.02 ± 0.01 | 0.02 ± 0.01 |
| | | P | 1.22 | 1.15 | 1.09 ± 0.01 | 0.99 ± 0.08 | 0.94 ± 0.1 | 0.83 ± 0.07 | 0.93 ± 0.17 |
| | | NP | 1.3 | 1.07 | 0.8 ± 0.11 | 0.12 ± 0.08 | 0.07 ± 0.01 | 0.08 ± 0.03 | 0.05 ± 0.04 |
| | | NPSi | 1.25 | 0.92 | 0.45 ± 0.24 | 0.38 ± 0.29 | 0.14 ± 0.01 | 0.13 ± 0.03 | 0.07 ± 0.07 |
| SiO2 | 2022 | Control | 2.63 ± 0.13 | 2.54 ± 0.08 | 2.05 ± 0.11 | 1.66 ± 0.14 | 1.72 ± 0.09 | 1.74 ± 0.05 | 1.84 ± 0.12 |
| | | N | 2.45 ± 0.17 | 2.28 ± 0.08 | 1.7 ± 0.21 | 0.9 ± 0.42 | 0.52 ± 0.53 | 0.17 ± 0.16 | 0.33 ± 0.41 |
| | | P | 1.98 ± 0.09 | 2.23 ± 0.03 | 1.47 ± 0.11 | 1.21 ± 0.12 | 1.14 ± 0.32 | 1.1 ± 0.61 | 1.77 ± 1.65 |
| | | NP | 3.41 ± 0.09 | 3.26 ± 0.13 | 2.3 ± 0.33 | 1.12 ± 0.15 | 1.22 ± 0.11 | 1.28 ± 0.07 | 1.17 ± 0.12 |
| | 2023 | Control | 2.13 | 1.9 | 1.41 ± 0.21 | 0.71 ± 0.22 | 0.07 ± 0.12 | 12.55 ± 0.21 | 1.08 ± 0.22 |
| | | N | 1.71 | 1.64 | 1.6 ± 0.1 | 1.61 ± 0.09 | 1.58 ± 0.17 | 1.38 ± 0.26 | 0.95 ± 0.19 |
| | | P | 2 | 2.28 | 1.95 ± 0.09 | 1.93 ± 0.07 | 1.73 ± 0.08 | 2.37 ± 0.19 | 1.11 ± 0.12 |
| | | NP | 2.88 | 2.98 | 2.6 ± 0.09 | 0.88 ± 0.18 | 0.83 ± 0.04 | 0.8 ± 0.14 | 1.11 ± 0.12 |
| | | NPSi | 25.02 | 25.41 | 8.66 ± 7.22 | 10.96 ± 10.11 | 10.91 ± 1.02 | 12.55 ± 0.91 | 14.98 ± 0.67 |

**Table S3.** Dissolved Organic Carbon (DOC) concentrations measured during the 7-day incubation experiments in 2022 and 2023. Values represent the mean ± standard deviation of replicate bottles (n = 2–3) for each treatment and day.

| | Treatment | Day 1 | Day 2 | Day 3 | Day 4 | Day 5 | Day 6 | Day 7 |
|---|---|---|---|---|---|---|---|---|
| **2022** | Control | 76.19 ± 3.86 | 72.94 | 108.08 ± 13.6 | 105.73 ± 15.08 | 119.18 ± 16.77 | 80.51 ± 3.3 | 100.89 ± 3.05 |
| | N | - | - | 88.26 ± 11.07 | 125.08 ± 25.25 | 129.38 ± 19.52 | 85.15 ± 6.6 | 118.54 ± 6.62 |
| | P | - | 73.8 | 77.72 ± 2.37 | 151.23 ± 19.12 | 83.23 ± 8.93 | 86.38 ± 6.45 | 112.64 ± 9.47 |
| | NP | - | - | 98.83 ± 17.44 | 145.03 ± 16.94 | 108.47 ± 7.3 | 139.58 ± 19.55 | 166.54 ± 8.87 |
| **2023** | Control | 65.45 ± 3.37 | 62.18 | 80.48 ± 11.87 | 86.1 ± 16.49 | 89.86 ± 10.75 | 135.44 ± 35 | 84.93 ± 6.96 |
| | N | - | 59.78 | 84.05 ± 5.43 | 75.33 ± 2.96 | 102.78 ± 3.5 | 94.53 ± 7.23 | 115.17 ± 34.61 |
| | P | - | 72.11 | 85.73 ± 4.71 | 98.02 ± 12.63 | 99.86 ± 22.86 | 102.1 ± 4.96 | - |
| | NP | - | 67.27 | 84.01 ± 7.23 | 96.67 ± 3.37 | 95.87 ± 4.51 | 101.19 ± 5.22 | 130.74 ± 20.36 |
| | NPSi | - | - | 82.61 ± 2.38 | 85.3 ± 9.77 | 135.63 ± 7.71 | 135.44 ± 11.42 | 123.75 ± 1.11 |

**Table S4. Carbonate System Parameters in the 2022 and 2023 Experiments.** Mean seawater carbonate system parameters across replicate measurements for each treatment and day. Dissolved inorganic carbon (DIC), partial pressure of carbon dioxide ($pCO_2$), bicarbonate ion ($HCO_3^-$), carbonate ion ($CO_3^{2-}$), and dissolved carbon dioxide ($CO_2$) were calculated using CO2SYS version 3.0 (Lewis et al. 1998), based on measured $pH_T$ (total scale) and total alkalinity (TA). Initial conditions were based on TA measured in control bottles (n = 3). Input uncertainties were derived by calculating the standard deviation of repeated Certified Reference Material (CRM) measurements (Dickson, 1990) and were used in CO2SYS to propagate analytical uncertainty. For TA and pH, ± values represent the standard deviation of replicate measurements (n=2-3). For all other parameters, ± values represent the root-mean-square (RMS) of propagated errors.

| Year | Treatment | Day | $pH_T$ | TA (µmol/kg) | DIC (µmol/kg) | $pCO_2$ (µatm) | $HCO_3^-$ (µmol/kg) | $CO_3^{2-}$ (µmol/kg) | $CO_2$ (µmol/kg) | ΩCa | ΩAr |
|---|---|---|---|---|---|---|---|---|---|---|---|
| 2022 | Control | 1 | 8.14 ± 0.001 | 2262.8 ± 3.6 | 1915.1 ± 7.8 | 354.3 ± 13.2 | 1664.6 ± 11.5 | 241.3 ± 4.7 | 9.2 ± 0.3 | 5.9 ± 0.3 | 3.9 ± 0.1 |
| | | 2 | 8.14 ± 0.006 | 2273 ± 7.9 | 1920.2 ± 7.7 | 334.6 ± 13 | 1666.8 ± 11.4 | 244.5 ± 4.7 | 8.9 ± 0.3 | 6 ± 0.3 | 4 ± 0.1 |
| | | 4 | 8.16 ± 0.006 | 2302.5 ± 25.3 | 1931.2 ± 7.9 | 308 ± 12.4 | 1666 ± 11.7 | 256.8 ± 4.8 | 8.4 ± 0.3 | 6.2 ± 0.3 | 4.1 ± 0.1 |
| | | 6 | 8.17 ± 0.006 | 2279.3 ± 2.8 | 1911 ± 8 | 326.3 ± 12.4 | 1647.7 ± 11.9 | 254.8 ± 4.9 | 8.5 ± 0.3 | 6.2 ± 0.3 | 4.1 ± 0.1 |
| | | 7 | 8.15 ± 0.006 | 2276.2 ± 1.3 | 1920.6 ± 7.9 | 345 ± 13 | 1664.7 ± 11.7 | 246.9 ± 4.8 | 9 ± 0.3 | 6 ± 0.3 | 4 ± 0.1 |
| | N | 2 | 8.14 ± 0.015 | 2276 ± 14 | 1924.8 ± 7.7 | 337.6 ± 13.1 | 1672.3 ± 11.4 | 243.5 ± 4.7 | 9 ± 0.4 | 5.9 ± 0.3 | 3.9 ± 0.1 |
| | | 4 | 8.16 ± 0.01 | 2293.9 ± 36.6 | 1925.5 ± 7.8 | 310.5 ± 12.5 | 1663 ± 11.6 | 254 ± 4.7 | 8.4 ± 0.3 | 6.2 ± 0.3 | 4.1 ± 0.1 |
| | | 6 | 8.18 ± 0.01 | 2287 ± 19 | 1908.9 ± 8.1 | 316.7 ± 12.1 | 1639.8 ± 12.1 | 260.9 ± 5 | 8.2 ± 0.3 | 6.4 ± 0.3 | 4.2 ± 0.1 |
| | | 7 | 8.06 ± 0.115 | 2293.1 ± 8.7 | 1985 ± 7.4 | 457.8 ± 17 | 1755.8 ± 10.7 | 217.4 ± 4.3 | 11.9 ± 0.4 | 5.3 ± 0.2 | 3.5 ± 0.1 |
| | P | 2 | 8.14 ± 0.006 | 2280.1 ± 28.8 | 1929.9 ± 7.7 | 340.1 ± 13.2 | 1677.7 ± 11.4 | 243.2 ± 4.7 | 9.1 ± 0.3 | 5.9 ± 0.3 | 3.9 ± 0.1 |
| | | 4 | 8.17 ± 0.006 | 2274 ± 9.2 | 1906.4 ± 7.8 | 302.5 ± 12.2 | 1644.3 ± 11.6 | 253.8 ± 4.7 | 8.2 ± 0.3 | 6.2 ± 0.3 | 4.1 ± 0.1 |
| | | 6 | 8.15 ± 0.01 | 2274.1 ± 1.5 | 1917.1 ± 7.9 | 341 ± 12.9 | 1660.5 ± 11.7 | 247.7 ± 4.8 | 8.9 ± 0.3 | 6 ± 0.3 | 4 ± 0.1 |
| | | 7 | 8.14 ± 0.006 | 2279.9 ± 3.7 | 1931.3 ± 7.8 | 358 ± 13.4 | 1679.8 ± 11.5 | 242.3 ± 4.7 | 9.3 ± 0.4 | 5.9 ± 0.3 | 3.9 ± 0.1 |
| | NP | 2 | 8.14 ± 0.001 | 2274.2 ± 4.8 | 1924.7 ± 7.7 | 337.9 ± 13.1 | 1672.8 ± 11.3 | 242.9 ± 4.6 | 9 ± 0.4 | 5.9 ± 0.3 | 3.9 ± 0.1 |
| | | 4 | 8.26 ± 0.053 | 2296.3 ± 11.9 | 1861.3 ± 8.5 | 233 ± 9.8 | 1556.6 ± 12.9 | 298.3 ± 5.4 | 6.3 ± 0.3 | 7.3 ± 0.3 | 4.8 ± 0.1 |
| | | 6 | 8.22 ± 0.127 | 2294 ± 5.5 | 1887.5 ± 8.4 | 296.1 ± 11.8 | 1599.6 ± 12.7 | 280.2 ± 5.3 | 7.7 ± 0.3 | 6.8 ± 0.3 | 4.6 ± 0.1 |
| | | 7 | 8.17 ± 0.183 | 2299.1 ± 17.4 | 1916.2 ± 8.2 | 353.7 ± 14.2 | 1642.5 ± 12.2 | 264.5 ± 5 | 9.2 ± 0.4 | 6.5 ± 0.3 | 4.3 ± 0.1 |
| 2023 | Control | 1 | 8.04 ± 0.006 | 2278.6 ± 1.6 | 1986.4 ± 5.6 | 493.6 ± 12.2 | 1766.9 ± 8.4 | 207.2 ± 3.6 | 12.3 ± 0.3 | 5.1 ± 0.2 | 3.4 ± 0.1 |
| | | 3 | 8.06 ± 0.007 | 2281.1 ± 1.9 | 1982.1 ± 5.7 | 476 ± 11.9 | 1758.5 ± 8.5 | 211.6 ± 3.6 | 11.9 ± 0.3 | 5.2 ± 0.2 | 3.5 ± 0.1 |
| | | 4 | 8.06 ± 0.007 | 2267 ± 1 | 1968.7 ± 6.4 | 465.3 ± 10.3 | 1746.5 ± 8.5 | 210.5 ± 3.4 | 11.8 ± 0.3 | 5.2 ± 0.2 | 3.4 ± 0.1 |
| | | 5 | 8.07 ± 0.007 | 2270.2 ± 7.9 | 1966.4 ± 6.4 | 442.3 ± 10 | 1741.1 ± 8.5 | 213.9 ± 3.4 | 11.4 ± 0.3 | 5.2 ± 0.2 | 3.5 ± 0.1 |
| | | 7 | 8.06 ± 0.012 | 2286 ± 36.2 | 1986.7 ± 6.4 | 451.8 ± 10.3 | 1763.4 ± 8.4 | 211.5 ± 3.3 | 11.7 ± 0.3 | 5.2 ± 0.2 | 3.4 ± 0.1 |
| | N | 3 | 8.05 ± 0.001 | 2282.6 ± 2.6 | 1986.9 ± 5.6 | 483.9 ± 12 | 1765.1 ± 8.4 | 209.6 ± 3.6 | 12.1 ± 0.3 | 5.1 ± 0.2 | 3.4 ± 0.1 |
| | | 4 | 8.06 (*) | 2267.6 ± * | 1965.2 ± 6.4 | 456.3 ± 10.1 | 1740.5 ± 8.6 | 213.1 ± 3.4 | 11.5 ± 0.3 | 5.2 ± 0.2 | 3.5 ± 0.1 |
| | | 5 | 8.06 ± 0.007 | 2269.7 ± 4.6 | 1967.1 ± 6.4 | 444.7 ± 10 | 1742.5 ± 8.5 | 213.1 ± 3.4 | 11.4 ± 0.3 | 5.2 ± 0.2 | 3.5 ± 0.1 |
| | | 7 | 8.04 ± 0.007 | 2273 ± 3.5 | 1983.5 ± 6.3 | 467.9 ± 10.6 | 1766.5 ± 8.3 | 204.9 ± 3.2 | 12.1 ± 0.3 | 5 ± 0.2 | 3.3 ± 0.1 |
| | P | 3 | 8.04 ± 0.001 | 2283 ± 6 | 1990.5 ± 5.6 | 486.7 ± 12.2 | 1771.3 ± 8.3 | 206.9 ± 3.5 | 12.3 ± 0.3 | 5.1 ± 0.2 | 3.4 ± 0.1 |
| | | 4 | 8.05 ± 0.001 | 2269.7 ± 2.1 | 1974.1 ± 6.4 | 485.3 ± 10.5 | 1752.9 ± 8.6 | 209.1 ± 3.4 | 12.1 ± 0.3 | 5.1 ± 0.2 | 3.4 ± 0.1 |
| | | 5 | 8.06 ± 0.007 | 2262.9 ± 3.9 | 1962.9 ± 6.4 | 447.7 ± 10.1 | 1740.3 ± 8.4 | 211.2 ± 3.3 | 11.5 ± 0.3 | 5.2 ± 0.2 | 3.4 ± 0.1 |
| | | 7 | 8.05 ± 0.001 | 2269.9 ± 6.6 | 1973.7 ± 6.3 | 451.8 ± 10.3 | 1753.1 ± 8.4 | 208.9 ± 3.3 | 11.7 ± 0.3 | 5.1 ± 0.2 | 3.4 ± 0.1 |
| | NP | 3 | 8.03 ± 0.001 | 2285.2 ± 6.8 | 1999.3 ± 5.5 | 510.7 ± 12.6 | 1783.8 ± 8.2 | 202.7 ± 3.5 | 12.8 ± 0.3 | 5 ± 0.2 | 3.3 ± 0.1 |
| | | 4 | 8.1 ± 0.007 | 2285.3 ± 4.3 | 1960.3 ± 6.6 | 430.4 ± 9.4 | 1722 ± 9 | 227.6 ± 3.6 | 10.7 ± 0.2 | 5.6 ± 0.2 | 3.7 ± 0.1 |
| | | 5 | 8.17 ± 0.001 | 2244.5 ± 54.6 | 1875.8 ± 6.9 | 321.1 ± 7.6 | 1614.3 ± 9.4 | 253.3 ± 3.8 | 8.3 ± 0.2 | 6.2 ± 0.3 | 4.1 ± 0.1 |
| | | 7 | 8.15 ± 0.007 | 2285.3 ± 7.3 | 1924.3 ± 6.8 | 340.7 ± 8.1 | 1665.4 ± 9.3 | 250.1 ± 3.8 | 8.8 ± 0.2 | 6.1 ± 0.3 | 4.1 ± 0.1 |
| | NPSi | 3 | 7.93 ± 0.001 | 2199.1 ± 7.7 | 1975.4 ± 4.8 | 641.4 ± 15.5 | 1797.8 ± 6.8 | 161.4 ± 2.8 | 16.2 ± 0.4 | 4 ± 0.2 | 2.6 ± 0.1 |
| | | 4 | 8.02 ± 0.028 | 2213.5 ± 16.2 | 1939.7 ± 6.2 | 515.3 ± 11 | 1733.9 ± 8.1 | 193 ± 3.2 | 12.8 ± 0.3 | 4.7 ± 0.2 | 3.2 ± 0.1 |
| | | 5 | 8.26 ± 0.014 | 2143.1 ± 38.3 | 1727.9 ± 7.1 | 234.5 ± 5.8 | 1444.2 ± 9.9 | 277.7 ± 4.1 | 6 ± 0.1 | 6.8 ± 0.3 | 4.5 ± 0.1 |
| | | 7 | 8.15 ± 0.021 | 2202.3 ± 5.2 | 1855.8 ± 6.7 | 335.6 ± 8 | 1609.7 ± 9 | 237.4 ± 3.6 | 8.7 ± 0.2 | 5.8 ± 0.2 | 3.9 ± 0.1 |

**Table S5.** Proportions of $HCO_3^-$, $CO_3^{2-}$, and $CO_2$ relative to the dissolved inorganic carbon (DIC) pool in each treatment and year. Colors represent lower (blue) and higher (orange) values than initial conditions (Control, Day 1) for each year. Bicarbonate ($HCO_3^-$) was dominant in all samples, ranging between 83-91%. Decreases in bicarbonate proportions corresponded with increases in $CO_3^{2-}$ during $CO_2$ photosynthetic uptake, especially after nutrient enrichment (NP and NPSi treatments).

| Year | Treatment | Day | % HCO3 | % CO3 | % CO2 |
|---|---|---|---|---|---|
| 2022 | Control | 1 | 86.9 | 12.6 | 0.5 |
| | | 2 | 86.8 | 12.7 | 0.5 |
| | | 4 | 86.3 | 13.3 | 0.4 |
| | | 6 | 86.2 | 13.3 | 0.4 |
| | | 7 | 86.7 | 12.9 | 0.5 |
| | N | 2 | 86.9 | 12.7 | 0.5 |
| | | 4 | 86.4 | 13.2 | 0.4 |
| | | 6 | 85.9 | 13.7 | 0.4 |
| | | 7 | 88.5 | 11.0 | 0.6 |
| | P | 2 | 86.9 | 12.6 | 0.5 |
| | | 4 | 86.3 | 13.3 | 0.4 |
| | | 6 | 86.6 | 12.9 | 0.5 |
| | | 7 | 87.0 | 12.5 | 0.5 |
| | NP | 2 | 86.9 | 12.6 | 0.5 |
| | | 4 | 83.6 | 16.0 | 0.3 |
| | | 6 | 84.7 | 14.8 | 0.4 |
| | | 7 | 85.7 | 13.8 | 0.5 |
| 2023 | Control | 1 | 88.9 | 10.4 | 0.6 |
| | | 3 | 88.7 | 10.7 | 0.6 |
| | | 4 | 88.7 | 10.7 | 0.6 |
| | | 5 | 88.5 | 10.9 | 0.6 |
| | | 7 | 88.8 | 10.6 | 0.6 |
| | N | 3 | 88.8 | 10.5 | 0.6 |
| | | 4 | 88.6 | 10.8 | 0.6 |
| | | 5 | 88.6 | 10.8 | 0.6 |
| | | 7 | 89.1 | 10.3 | 0.6 |
| | P | 3 | 89.0 | 10.4 | 0.6 |
| | | 4 | 88.8 | 10.6 | 0.6 |
| | | 5 | 88.7 | 10.8 | 0.6 |
| | | 7 | 88.8 | 10.6 | 0.6 |
| | NP | 3 | 89.2 | 10.1 | 0.6 |
| | | 4 | 87.8 | 11.6 | 0.5 |
| | | 5 | 86.1 | 13.5 | 0.4 |
| | | 7 | 86.5 | 13.0 | 0.5 |
| | NPSi | 3 | 91.0 | 8.2 | 0.8 |
| | | 4 | 89.4 | 9.9 | 0.7 |
| | | 5 | 83.6 | 16.1 | 0.3 |
| | | 7 | 86.7 | 12.8 | 0.5 |

**Table S6. Size-fractionated chlorophyll *a* concentration (µg $L^{-1}$) from surface water at different locations around Ishigaki Island. Reference dissolved inorganic nutrient concentrations at other locations around Ishigaki Island in 2023 (n=1)**. The first two rows show initial conditions from the incubation experiments in 2022 and 2023. If the N:P ratio is lower than 16, nitrogen is limiting relative to phosphorus, but if it exceeds 16, there is excess N. If Si:N is lower than 1, silica is limiting relative to nitrogen, but if it is higher than 1, there is excess silica, making conditions favorable for diatoms (given that nitrogen is still available). If Si:P is lower than 16, silica is limiting relative to phosphorus. If it is higher than 16, there is excess silica.

| Study Site | Coordinates | Environment | Total Chl *a* | 0.2 – 1 µm | 1 - 10 µm | 10 - 150 µm | % 0.2 – 1 µm | % 1 - 10 µm | % 10 - 150 µm | $NO_2^-$ | $NH_4^+$ | $NO_3^-$ | DIN | $PO_4^{3-}$ | $SiO_2$ | N:P | Si:N | Si:P |
|---|---|---|---|---|---|---|---|---|---|---|---|---|---|---|---|---|---|---|
| **Experiment 2022** | 24°21'52"N 124°15'12"E | Reef | 0.55 | - | - | - | - | - | - | 0.15 | 1.24 | 1.18 | 2.56 | 0.23 | 2.63 | 11.53 | 1.04 | 11.63 |
| **Experiment 2023** | 24°21'52"N 124°15'12"E | *Reef* | 0.64 | 0.04 | 0.1 | 0.5 | 6.3 | 15.6 | 78.1 | 0.05 | 0.20 | 0.55 | 0.80 | 0.07 | 2.13 | 11.63 | 2.67 | 31.10 |
| **Shiraho Reef Crest** | 24°23'03"N 124°15'26"E | *Reef* | 0.38 | 0.15 | 0.04 | 0.18 | 39.5 | 10.5 | 50.0 | 0.11 | 0.22 | 0.79 | 1.12 | 0.19 | 1.24 | 6.00 | 1.10 | 6.62 |
| **Shiraho Reef** | 24°21'11"N 124°15'04"E | *Reef* | 0.61 | 0.15 | 0.07 | 0.38 | 24.6 | 11.5 | 63.9 | 0.11 | 0.25 | 0.42 | 0.79 | 0.13 | 1.08 | 6.14 | 1.36 | 8.37 |
| **Fukido Reef** | 24°29'53"N 124°13'31"E | *Reef* | 0.58 | 0.21 | 0.13 | 0.24 | 36.2 | 22.4 | 41.4 | 0.13 | 0.27 | 0.67 | 1.07 | 0.12 | 2.28 | 8.75 | 2.13 | 18.64 |
| **Todoroki River Mouth** | **24°23'00"N 124°15'10"E** | ***River*** | **2.15** | **0.11** | **0.19** | **1.82** | **5.1** | 8.8 | **86.0** | **0.26** | **0.51** | **9.48** | **10.25** | **1.01** | **33.85** | **10.19** | **3.30** | **33.67** |
| **Nagura Bay** | 24°24'07"N 124°08'19"E | *River* | 1.97 | 0.60 | 0.15 | 1.05 | 30.5 | 7.6 | 61.9 | 0.08 | 0.26 | 0.33 | 0.67 | 0.14 | 4.08 | 4.82 | 6.11 | 29.45 |
| **Fukido River Mouth** | 24°29'27"N 124°13'42"E | *River* | 1.41 | 0.53 | 0.22 | 0.66 | 37.6 | 15.6 | 46.8 | 0.09 | 0.37 | 0.39 | 0.86 | 0.12 | 2.59 | 7.35 | 3.03 | 22.24 |
| **Miyara River** | 24°21'11"N 124°12'56"E | *River* | 1.26 | 0.41 | 0.23 | 0.58 | 32.5 | 18.3 | 49.2 | 0.12 | 0.19 | 0.70 | 1.01 | 0.15 | 3.08 | 6.91 | 3.06 | 21.16 |

**Statistical analyses**

*Total Chlorophyll a*

Given the high variability among chlorophyll concentration results resulting from different nutrient enrichment regimes, total and size-fractionated Chl *a* values did not follow a normal distribution. Therefore, Chl *a* data were log-transformed to perform statistical analyses. To evaluate significant differences between days and between treatments, a Linear Mixed Effects Model was fitted separately for each year (2022: R2= 0.967; 2023: R2= 0.948. The fixed effects included treatment, day, and their interaction. Replicate was treated as a random effect to account for repeated measures. Post hoc pairwise comparisons of estimated marginal means (EMMs) were performed using Tukey-adjusted contrasts. These comparisons evaluated (1) differences between days within each treatment (Table S7) and differences between treatments within each sampling day (Table S8).

**Table S7. Post hoc pairwise comparisons of total Chlorophyll a concentrations between days within each treatment for 2022 and 2023. Significant differences (Tukey-adjusted p < 0.05) are shown in bold. Light gray values denote non-significant differences.**

| | Treatment | Day 1 - 2 | Day 2 - 3 | Day 3 - 4 | Day 4 - 5 | Day 5 - 6 | Day 6 - 7 |
|---|---|---|---|---|---|---|---|
| **2022** | Control | 1 | **<0.001** | 0.263 | **<0.001** | 0.892 | 0.702 |
| | N | 0.347 | **<0.001** | 0.99 | 0.869 | 1 | 1 |
| | P | 0.477 | **<0.001** | **<0.001** | **<0.001** | 0.96 | 0.976 |
| | NP | **0.006** | **<0.001** | **<0.001** | **<0.001** | **<0.001** | 0.894 |
| **2023** | Control | 0.995 | 0.996 | 1 | 0.48 | 0.999 | 0.415 |
| | N | 1 | 0.983 | 1 | 0.176 | 0.999 | 0.173 |
| | P | 0.792 | 0.999 | 0.626 | 0.729 | 1 | 0.085 |
| | NP | 1 | **<0.001** | **<0.001** | **0.014** | 0.149 | 0.084 |
| | NPSi | 1 | **<0.001** | **<0.001** | 0.164 | **<0.001** | 0.91 |

**Table S8. Post hoc pairwise comparisons of total Chlorophyll a concentrations between treatments within each day for 2022 and 2023. Significant differences (Tukey-adjusted p < 0.05) are shown in bold. Light gray values denote non-significant differences.**

| Year | Day | Control - N | Control - P | Control - NP | N - P | N - NP | NP - P | Control - NPSI | N - NPSI | NP - NPSI |
|---|---|---|---|---|---|---|---|---|---|---|
| 2022 | 1 | 1 | 1 | 1 | 1 | 1 | 1 | | | |
| | 2 | 0.268 | 0.349 | **0.016** | 0.998 | 0.54 | 0.438 | | | |
| | 3 | 0.813 | 0.113 | **<0.001** | 0.481 | **0.004** | 0.119 | | | |
| | 4 | **0.013** | 0.994 | **<0.001** | **0.007** | **<0.001** | **<0.001** | | | |
| | 5 | **<0.001** | 0.82 | **<0.001** | **<0.001** | **<0.001** | **<0.001** | | | |
| | 6 | **<0.001** | 0.903 | **<0.001** | **<0.001** | **<0.001** | **<0.001** | | | |
| | 7 | **<0.001** | 0.607 | **<0.001** | **<0.001** | **0.004** | **<0.001** | | | |
| 2023 | 1 | 1 | 1 | 1 | 1 | 1 | 1 | 1 | 1 | 1 |
| | 2 | 0.978 | 0.961 | 0.999 | 0.719 | 0.998 | 0.878 | 0.992 | 1 | 1 |
| | 3 | 0.945 | 0.483 | **<0.001** | 0.139 | **0.002** | **<0.001** | **<0.001** | **0.001** | 1 |
| | 4 | 0.943 | 0.999 | **<0.001** | 0.86 | **<0.001** | **<0.001** | **<0.001** | **<0.001** | **0.029** |
| | 5 | 0.711 | 0.979 | **<0.001** | 0.363 | **<0.001** | **<0.001** | **<0.001** | **<0.001** | **0.003** |
| | 6 | 0.24 | 0.989 | **<0.001** | 0.092 | **<0.001** | **<0.001** | **<0.001** | **<0.001** | 0.182 |
| | 7 | 0.101 | 0.986 | **<0.001** | 0.395 | **<0.001** | **<0.001** | **<0.001** | **0.001** | 0.732 |

*Size-fractionated Chl* a (2023)

To evaluate how nutrient treatments, experimental day, and phytoplankton size classes influenced chlorophyll *a* concentrations in 2023, a linear mixed-effects model (LMM) including random effects was fitted (R2 = 0.938). This model was selected due to the hierarchical nature of the data, with repeated measures from the same replicate units (bottles). The three-way interaction between treatment, day, and filter was statistically significant, indicating that nutrient addition influenced not only the Chl-a concentration but also the overall size structure of phytoplankton over time.

**Table S9.** Pairwise comparisons of variation in Chl-a concentrations between consecutive days within each treatment and size class. Significant differences (Tukey-adjusted $p < 0.05$) are shown in bold. Light gray values denote non-significant differences.

| Treatment | Size class | Day 1 - 2 | Day 2 - 3 | Day 3 - 4 | Day 4 - 5 | Day 5 - 6 | Day 6 - 7 |
|---|---|---|---|---|---|---|---|
| Control | Pico | 0.101 | 0.43 | 1 | 0.852 | 0.836 | 0.999 |
| N | | **<0.001** | 1 | 1 | 0.71 | 1 | 0.998 |
| P | | **0.001** | 1 | 0.977 | 0.829 | 0.938 | 1 |
| NP | | **0.003** | **<0.001** | **0.002** | 0.979 | 0.892 | 0.798 |
| NPSi | | **0.01** | **<0.001** | 0.073 | 0.771 | 0.244 | 0.502 |
| Control | Nano | 0.994 | 0.956 | 0.975 | 0.999 | 1 | 1 |
| N | | 0.999 | 1 | 0.983 | 0.986 | 1 | 0.994 |
| P | | 0.996 | 0.998 | 0.999 | 1 | 0.993 | 1 |
| NP | | 1 | 0.161 | **0.003** | 0.122 | 0.991 | 0.828 |
| NPSi | | 0.999 | 0.355 | **<0.001** | 0.064 | 0.241 | 0.608 |
| Control | Micro | 0.974 | 1 | 1 | 0.809 | 1 | 0.543 |
| N | | 0.911 | 0.958 | 1 | 0.372 | 1 | 0.305 |
| P | | 0.425 | 1 | 0.829 | 0.93 | 1 | 0.136 |
| NP | | 0.96 | **0.001** | **<0.001** | 0.142 | 0.126 | 0.306 |
| NPSi | | 0.984 | **0.001** | **<0.001** | 0.603 | **0.034** | 0.997 |

**Table S10.** Pairwise comparisons of treatment-level differences in Chl *a* concentration within each size class and sampling day. Significant differences (Tukey-adjusted $p < 0.05$), shown in bold, indicate different Chl-a values on a given day and size fraction. Light gray values denote non-significant differences.

| Day | Size class | **Control - N** | **Control - P** | **Control - NP** | **N - P** | **N - NP** | **NP - P** | **Control - NPSi** | **N - NPSi** | **NPSi - P** | **NP - NPSi** |
|---|---|---|---|---|---|---|---|---|---|---|---|
| 1 | Pico | 1 | 1 | 1 | 1 | 1 | 1 | 1 | 1 | 1 | 1 |
| 2 | | 0.268 | 0.743 | 0.818 | 0.932 | 0.884 | 1 | 0.948 | 0.708 | 0.989 | 0.997 |
| 3 | | 0.993 | 0.97 | **0.005** | 0.832 | **0.02** | **<0.001** | **0.023** | 0.073 | **0.003** | 0.989 |
| 4 | | 0.997 | 1 | **<0.001** | 0.998 | **<0.001** | **<0.001** | **<0.001** | **<0.001** | **<0.001** | 0.562 |
| 5 | | 0.976 | 1 | **<0.001** | 0.986 | **<0.001** | **<0.001** | **0.027** | 0.122 | **0.033** | 0.245 |
| 6 | | 0.25 | 0.998 | **<0.001** | 0.422 | **<0.001** | **<0.001** | 0.228 | 1 | 0.392 | **<0.001** |
| 7 | | **0.03** | 0.988 | **<0.001** | 0.193 | **<0.001** | **<0.001** | 0.982 | 0.215 | 1 | **<0.001** |
| 1 | Nano | 1 | 1 | 1 | 1 | 1 | 1 | 1 | 1 | 1 | 1 |
| 2 | | 0.845 | 1 | 0.964 | 0.862 | 0.996 | 0.97 | 0.823 | 1 | 0.841 | 0.994 |
| 3 | | 0.993 | 0.651 | 0.248 | 0.387 | 0.48 | **0.007** | 0.234 | 0.461 | **0.007** | 1 |
| 4 | | 0.989 | 1 | **<0.001** | 0.975 | **<0.001** | **<0.001** | **<0.001** | **<0.001** | **<0.001** | 0.303 |
| 5 | | 0.936 | 1 | **0.005** | 0.874 | 0.052 | **0.003** | **<0.001** | **<0.001** | **<0.001** | 0.436 |
| 6 | | 0.947 | 0.932 | **0.038** | 0.542 | 0.214 | **0.003** | **0.018** | 0.127 | **0.001** | 0.999 |
| 7 | | 0.809 | 0.874 | **0.002** | 0.297 | 0.052 | **<0.001** | 0.85 | 1 | 0.371 | 0.102 |
| 1 | Micro | 1 | 1 | 1 | 1 | 1 | 1 | 1 | 1 | 1 | 1 |
| 2 | | 0.999 | 0.853 | 1 | 0.941 | 1 | 0.886 | 1 | 0.998 | 0.818 | 1 |
| 3 | | 0.958 | 0.673 | **0.003** | 0.263 | **0.026** | **<0.001** | **0.002** | **0.016** | **<0.001** | 1 |
| 4 | | 0.989 | 0.998 | **<0.001** | 0.932 | **<0.001** | **<0.001** | **<0.001** | **<0.001** | **<0.001** | 0.113 |
| 5 | | 0.804 | 0.977 | **<0.001** | 0.444 | **<0.001** | **<0.001** | **<0.001** | **<0.001** | **<0.001** | **0.014** |
| 6 | | 0.63 | 0.987 | **<0.001** | 0.323 | **0.002** | **<0.001** | **<0.001** | **<0.001** | **<0.001** | 0.051 |
| 7 | | 0.41 | 0.991 | **<0.001** | 0.789 | **0.002** | **<0.001** | **<0.001** | **0.003** | **<0.001** | 1 |

*Dissolved Inorganic Nutrients*

**Table S11. Differences between days, within treatments**

| Nutrient | Year | Treatment | Day1 - Day2 | Day2 - Day3 | Day3 - Day4 | Day4 - Day5 | Day5 - Day6 | Day6 - Day7 |
|---|---|---|---|---|---|---|---|---|
| DIN | 2022 ($R^2$ = 0.978) | Control | 1 | 1 | **0.002** | 0.939 | 0.99 | 0.964 |
| | | N | 1 | 1 | 0.997 | 0.99 | 0.781 | 1 |
| | | P | 1 | 1 | 0.859 | **<0.001** | 0.208 | 0.183 |
| | | NP | 0.991 | 0.97 | **<0.001** | **<0.001** | 0.898 | 0.466 |
| | 2023 ($R^2$ = 0.981) | Control | 0.968 | 0.997 | 1 | 0.999 | 1 | 0.966 |
| | | N | 1 | 1 | 1 | 0.995 | 1 | 0.998 |
| | | P | 0.912 | 1 | 1 | 0.997 | 1 | 1 |
| | | NP | 1 | 0.963 | **<0.001** | **<0.001** | 1 | 0.872 |
| | | NPSi | 0.948 | 0.806 | **<0.001** | **0.009** | 0.564 | 0.819 |
| $NH_4^+$ | 2022 ($R^2$ = 0.614) | Control | 1 | 0.371 | **0.041** | 0.97 | 0.987 | 0.992 |
| | | N | 0.874 | 1 | 1 | 0.201 | 1 | 0.97 |
| | | P | 1 | 0.414 | 0.926 | **<0.001** | 0.259 | 0.095 |
| | | NP | **0.049** | 0.363 | 1 | 0.723 | 0.976 | 0.542 |
| | 2023 ($R^2$ = 0.644) | Control | 0.999 | 1 | 0.994 | 1 | 1 | 0.979 |
| | | N | 1 | 0.959 | 0.875 | 0.89 | 1 | 1 |
| | | P | 0.999 | 0.998 | 1 | 0.937 | 1 | 1 |
| | | NP | 0.943 | 0.999 | **0.049** | 0.264 | 1 | 0.496 |
| | | NPSi | 0.998 | 0.545 | 0.925 | **0.006** | 0.746 | 0.943 |
| $NO_2^-$ | 2022 ($R^2$ = 0.848) | Control | 0.999 | **<0.001** | 0.61 | 1 | 0.971 | 1 |
| | | N | **0.018** | 0.346 | **0.006** | 1 | 0.326 | 0.053 |
| | | P | 1 | **0.005** | 0.99 | 0.848 | 0.991 | 1 |
| | | NP | 0.331 | **0.001** | **<0.001** | **0.011** | 0.998 | 1 |
| | 2023 | Control | 0.848 | 0.11 | 0.166 | 0.997 | 0.995 | 0.983 |

| | | | | | | | | |
|---|---|---|---|---|---|---|---|---|
| | ($R^2$ = 0.956) | N | **0.001** | 0.565 | **<0.001** | 0.998 | 0.931 | 0.693 |
| | | P | **0.046** | 0.999 | 1 | 1 | 0.965 | 0.984 |
| | | NP | 0.895 | **0.014** | 0.912 | **<0.001** | 0.773 | 0.926 |
| | | NPSi | 0.167 | **0.01** | **<0.001** | **<0.001** | 1 | 0.53 |
| $NO_3^-$ | 2022 ($R^2$ = 0.996) | Control | 0.334 | **<0.001** | 0.796 | 1 | 0.999 | 1 |
| | | N | 1 | 0.994 | 0.949 | 0.991 | 0.094 | 0.995 |
| | | P | 0.953 | **<0.001** | 1 | 0.998 | 1 | 0.942 |
| | | NP | 0.995 | 0.222 | **<0.001** | **<0.001** | 0.996 | 0.996 |
| | 2023 ($R^2$ = 0.991) | Control | 0.96 | 0.928 | 1 | 1 | 1 | 1 |
| | | N | 1 | 1 | 1 | 0.987 | 1 | 0.989 |
| | | P | 0.879 | 1 | 1 | 1 | 1 | 1 |
| | | NP | 0.998 | 0.869 | **<0.001** | **<0.001** | 0.999 | 0.997 |
| | | NPSi | 0.881 | 0.291 | **<0.001** | 1 | 0.994 | 0.998 |
| $PO_4^{3-}$ | 2022 ($R^2$ = 0.969) | Control | 0.453 | 0.965 | 0.474 | 0.963 | 0.991 | 1 |
| | | N | 0.332 | 0.817 | 1 | 1 | **0.02** | 1 |
| | | P | 0.968 | 0.145 | 0.155 | 0.952 | 1 | 0.99 |
| | | NP | 0.939 | 0.747 | **<0.001** | 0.129 | 0.622 | 1 |
| | 2023 ($R^2$ = 0.948) | Control | 1 | 1 | 1 | 1 | 1 | 1 |
| | | N | 1 | 1 | 1 | 1 | 1 | 1 |
| | | P | 1 | 1 | 0.958 | 0.995 | 0.897 | 0.94 |
| | | NP | 0.909 | 0.508 | **<0.001** | 0.965 | 1 | 0.993 |
| | | NPSi | 0.262 | **0.019** | 0.928 | **0.011** | 1 | 0.997 |
| $SiO_2$ | 2022 ($R^2$ = 0.762) | Control | 1 | 0.892 | 0.932 | 1 | 1 | 1 |
| | | N | 1 | 0.704 | 0.07 | 0.44 | 0.569 | 0.984 |
| | | P | 0.995 | 0.352 | 0.974 | 1 | 1 | 0.791 |
| | | NP | 1 | 0.39 | **0.017** | 1 | 1 | 1 |
| | 2023 | Control | 1 | 0.999 | 0.864 | 0.889 | 0.961 | 0.901 |

| | | | | | | | | |
|---|---|---|---|---|---|---|---|---|
| | ($R^2$ = 0.768) | N | 1 | 1 | 1 | 1 | 1 | 0.992 |
| | | P | 1 | 1 | 1 | 1 | 0.99 | 0.783 |
| | | NP | 1 | 1 | 0.295 | 1 | 1 | 0.998 |
| | | NPSi | 0.918 | 0.105 | 0.994 | 0.742 | 0.999 | 0.977 |

**Table S12. Differences between treatments**

| Nutrient | Year | Day | Control - N | Control - P | Control - NP | N - P | N - NP | NP - P | Control - NPSi | N - NPSi | NPSi - P | NP - NPSi |
|---|---|---|---|---|---|---|---|---|---|---|---|---|
| DIN | 2022 | 1 | **<0.001** | 0.593 | **<0.001** | **<0.001** | 0.998 | **<0.001** | | | | |
| | | 2 | **<0.001** | 0.443 | **<0.001** | **<0.001** | 0.927 | **<0.001** | | | | |
| | | 3 | **<0.001** | 0.747 | **<0.001** | **<0.001** | 0.591 | **<0.001** | | | | |
| | | 4 | **<0.001** | **<0.001** | **<0.001** | **<0.001** | **<0.001** | 0.567 | | | | |
| | | 5 | **<0.001** | 0.342 | 0.196 | **<0.001** | **<0.001** | 0.955 | | | | |
| | | 6 | **<0.001** | **0.007** | **0.03** | **<0.001** | **<0.001** | 0.954 | | | | |
| | | 7 | **<0.001** | 0.274 | 0.257 | **<0.001** | **<0.001** | 1 | | | | |
| | 2023 | 1 | **<0.001** | 0.81 | **<0.001** | **<0.001** | 1 | **<0.001** | **<0.001** | 1 | **<0.001** | 1 |
| | | 2 | **<0.001** | 0.907 | **<0.001** | **<0.001** | 0.998 | **<0.001** | **<0.001** | 0.808 | **<0.001** | 0.927 |
| | | 3 | **<0.001** | 0.124 | **<0.001** | **<0.001** | 0.415 | **<0.001** | **<0.001** | **0.004** | **<0.001** | 0.25 |
| | | 4 | **<0.001** | 0.175 | **<0.001** | **<0.001** | **<0.001** | **<0.001** | **0.003** | **<0.001** | 0.36 | **<0.001** |
| | | 5 | **<0.001** | 0.11 | **0.003** | **<0.001** | **<0.001** | 0.661 | 0.768 | **<0.001** | 0.678 | 0.074 |
| | | 6 | **<0.001** | 0.062 | **0.005** | **<0.001** | **<0.001** | 0.871 | 0.992 | **<0.001** | **0.021** | **0.001** |
| | | 7 | **<0.001** | 0.548 | 0.699 | **<0.001** | **<0.001** | 0.995 | 1 | **<0.001** | 0.575 | 0.726 |
| $NH_4^+$ | 2022 | 1 | 0.922 | 0.96 | 0.05 | 0.999 | 0.194 | 0.149 | | | | |
| | | 2 | 0.476 | 0.981 | 0.846 | 0.712 | 0.119 | 0.63 | | | | |
| | | 3 | 0.978 | 0.99 | 0.853 | 0.893 | 0.978 | 0.684 | | | | |
| | | 4 | 0.072 | **<0.001** | 0.063 | 0.194 | 1 | 0.217 | | | | |
| | | 5 | 0.767 | 1 | 0.417 | 0.826 | 0.902 | 0.475 | | | | |
| | | 6 | 1 | 0.365 | 0.221 | 0.42 | 0.263 | 0.99 | | | | |

| | | | | | | | | | | | | |
|---|---|---|---|---|---|---|---|---|---|---|---|---|
| | | 7 | 1 | 0.971 | 0.836 | 0.986 | 0.782 | 0.578 | | | | |
| | 2023 | 1 | 0.979 | 0.998 | 1 | 0.899 | 0.971 | 0.999 | 0.856 | 0.53 | 0.962 | 0.881 |
| | | 2 | 1 | 0.997 | 0.54 | 0.989 | 0.448 | 0.742 | 0.901 | 0.838 | 0.981 | 0.964 |
| | | 3 | 0.971 | 0.937 | **0.046** | 0.999 | 0.177 | 0.396 | **0.006** | **0.03** | 0.114 | 0.934 |
| | | 4 | 0.867 | 0.995 | **<0.001** | 0.966 | **0.002** | **<0.001** | **0.008** | **0.043** | **0.008** | 0.751 |
| | | 5 | 0.995 | 1 | **0.012** | 0.986 | **0.032** | **0.008** | 0.975 | 0.999 | 0.95 | 0.055 |
| | | 6 | 0.83 | 0.998 | **0.004** | 0.944 | 0.06 | **0.009** | 0.98 | 0.499 | 0.909 | **<0.001** |
| | | 7 | 0.99 | 1 | 0.884 | 0.966 | 0.985 | 0.802 | 0.999 | 0.935 | 1 | 0.729 |
| $NO_2^-$ | 2022 | 1 | **0.005** | 0.264 | **<0.001** | 0.274 | 0.242 | **0.004** | | | | |
| | | 2 | 0.915 | 0.521 | **0.007** | 0.883 | **0.032** | 0.148 | | | | |
| | | 3 | **<0.001** | 0.781 | **<0.001** | **<0.001** | 0.622 | **0.001** | | | | |
| | | 4 | **<0.001** | **0.039** | **0.036** | 0.252 | 0.267 | 1 | | | | |
| | | 5 | **<0.001** | 0.292 | 0.987 | **0.023** | **<0.001** | 0.209 | | | | |
| | | 6 | **<0.001** | **0.024** | 0.999 | **0.001** | **<0.001** | **0.018** | | | | |
| | | 7 | **<0.001** | **0.012** | 1 | **<0.001** | **<0.001** | **0.015** | | | | |
| | 2023 | 1 | 0.203 | **0.003** | 0.977 | 0.467 | 0.499 | **0.017** | 0.989 | 0.436 | **0.013** | 1 |
| | | 2 | **<0.001** | 0.994 | 0.991 | **<0.001** | **<0.001** | 0.909 | 0.466 | **0.009** | 0.253 | 0.744 |
| | | 3 | **<0.001** | 0.344 | **<0.001** | **<0.001** | 0.774 | **<0.001** | **<0.001** | 0.886 | **<0.001** | 0.254 |
| | | 4 | **<0.001** | **0.005** | **<0.001** | **<0.001** | **0.003** | **<0.001** | **<0.001** | **<0.001** | 0.179 | **<0.001** |
| | | 5 | **<0.001** | **0.013** | 0.997 | **<0.001** | **<0.001** | **0.005** | 0.999 | **<0.001** | **0.007** | 1 |
| | | 6 | **<0.001** | **0.022** | 0.864 | **<0.001** | **<0.001** | **0.002** | 0.996 | **<0.001** | 0.053 | 0.663 |
| | | 7 | **<0.001** | 0.063 | 0.939 | **<0.001** | **<0.001** | **0.006** | 0.796 | **<0.001** | 0.478 | 0.301 |
| $NO_3^-$ | 2022 | 1 | **<0.001** | **0.045** | **<0.001** | **<0.001** | 0.996 | **<0.001** | | | | |
| | | 2 | **<0.001** | **0.002** | **<0.001** | **<0.001** | 0.925 | **<0.001** | | | | |
| | | 3 | **<0.001** | **0.009** | **<0.001** | **<0.001** | 0.111 | **<0.001** | | | | |
| | | 4 | **<0.001** | **<0.001** | **<0.001** | **<0.001** | **<0.001** | **<0.001** | | | | |
| | | 5 | **<0.001** | **<0.001** | 0.933 | **<0.001** | **<0.001** | **0.015** | | | | |

| | | | | | | | | | | | | |
|---|---|---|---|---|---|---|---|---|---|---|---|---|
| | | 6 | **<0.001** | **<0.001** | 0.272 | **<0.001** | **<0.001** | **0.033** | | | | |
| | | 7 | **<0.001** | **<0.001** | 0.096 | **<0.001** | **<0.001** | **0.011** | | | | |
| | 2023 | 1 | **<0.001** | 0.748 | **<0.001** | **<0.001** | 1 | **<0.001** | **<0.001** | 0.998 | **<0.001** | 1 |
| | | 2 | **<0.001** | 0.877 | **<0.001** | **<0.001** | 0.98 | **<0.001** | **<0.001** | 0.587 | **<0.001** | 0.894 |
| | | 3 | **<0.001** | 0.059 | **<0.001** | **<0.001** | 0.082 | **<0.001** | **<0.001** | **<0.001** | **<0.001** | **0.029** |
| | | 4 | **<0.001** | **0.035** | **<0.001** | **<0.001** | **<0.001** | **<0.001** | 0.844 | **<0.001** | 0.193 | **<0.001** |
| | | 5 | **<0.001** | **0.004** | 0.678 | **<0.001** | **<0.001** | 0.127 | 0.836 | **<0.001** | 0.068 | 0.999 |
| | | 6 | **<0.001** | **0.007** | 0.968 | **<0.001** | **<0.001** | **0.04** | 1 | **<0.001** | **0.011** | 0.988 |
| | | 7 | **<0.001** | 0.103 | 0.9 | **<0.001** | **<0.001** | 0.334 | 0.994 | **<0.001** | 0.227 | 0.991 |
| $PO_4^{3-}$ | 2022 | 1 | 0.91 | **<0.001** | **<0.001** | **<0.001** | **<0.001** | 0.79 | | | | |
| | | 2 | 0.967 | **<0.001** | **<0.001** | **<0.001** | **<0.001** | 0.77 | | | | |
| | | 3 | **0.048** | **<0.001** | **<0.001** | **<0.001** | **<0.001** | 0.999 | | | | |
| | | 4 | 0.879 | **<0.001** | **0.026** | **<0.001** | **0.003** | **<0.001** | | | | |
| | | 5 | 0.509 | **<0.001** | 0.858 | **<0.001** | 0.969 | **<0.001** | | | | |
| | | 6 | 0.621 | **<0.001** | 0.994 | **<0.001** | 0.778 | **<0.001** | | | | |
| | | 7 | 0.833 | **<0.001** | 0.973 | **<0.001** | 0.581 | **<0.001** | | | | |
| | 2023 | 1 | 0.986 | **<0.001** | **<0.001** | **<0.001** | **<0.001** | 0.997 | **<0.001** | **<0.001** | 0.999 | 1 |
| | | 2 | 0.982 | **<0.001** | **<0.001** | **<0.001** | **<0.001** | 0.996 | **<0.001** | **<0.001** | 0.489 | 0.717 |
| | | 3 | 0.971 | **<0.001** | **<0.001** | **<0.001** | **<0.001** | 0.096 | **<0.001** | **<0.001** | **<0.001** | **0.001** |
| | | 4 | 0.896 | **<0.001** | 0.982 | **<0.001** | 0.505 | **<0.001** | **0.003** | **<0.001** | **<0.001** | **0.005** |
| | | 5 | 0.903 | **<0.001** | 1 | **<0.001** | 0.938 | **<0.001** | 0.757 | 0.254 | **<0.001** | 0.694 |
| | | 6 | 0.965 | **<0.001** | 0.99 | **<0.001** | 0.794 | **<0.001** | 0.727 | 0.336 | **<0.001** | 0.936 |
| | | 7 | 0.922 | **<0.001** | 0.995 | **<0.001** | 0.988 | **<0.001** | 0.998 | 0.781 | **<0.001** | 0.954 |
| $SiO_2$ | 2022 | 1 | 0.986 | 0.548 | 0.553 | 0.755 | 0.356 | 0.053 | | | | |
| | | 2 | 0.956 | 0.922 | 0.594 | 0.999 | 0.303 | 0.25 | | | | |
| | | 3 | 0.841 | 0.49 | 0.955 | 0.931 | 0.541 | 0.227 | | | | |
| | | 4 | 0.09 | 0.595 | 0.42 | 0.655 | 0.82 | 0.991 | | | | |

| | | | | | | | | | | | | |
|---|---|---|---|---|---|---|---|---|---|---|---|---|
| | | 5 | **<0.001** | 0.339 | 0.637 | 0.066 | **0.048** | 0.984 | | | | |
| | | 6 | **<0.001** | 0.19 | 0.595 | **0.003** | **<0.001** | 0.859 | | | | |
| | | 7 | **<0.001** | 0.704 | 0.277 | **<0.001** | **0.005** | 0.878 | | | | |
| | 2023 | 1 | 0.998 | 1 | 0.994 | 1 | 0.956 | 0.988 | **<0.001** | **<0.001** | **<0.001** | **0.001** |
| | | 2 | 1 | 0.999 | 0.979 | 0.997 | 0.958 | 0.997 | **0.008** | **0.006** | **0.015** | **0.036** |
| | | 3 | 0.999 | 0.979 | 0.743 | 0.996 | 0.859 | 0.984 | **0.045** | 0.075 | 0.257 | 0.447 |
| | | 4 | 0.659 | 0.469 | 0.991 | 0.997 | 0.856 | 0.673 | **<0.001** | **0.02** | **0.046** | **0.001** |
| | | 5 | 0.082 | 0.056 | 0.491 | 1 | 0.843 | 0.754 | **<0.001** | **<0.001** | **<0.001** | **<0.001** |
| | | 6 | 0.522 | 0.085 | 0.943 | 0.828 | 0.921 | 0.345 | **<0.001** | **<0.001** | **0.002** | **<0.001** |
| | | 7 | 1 | 1 | 1 | 1 | 0.999 | 1 | **<0.001** | **<0.001** | **<0.001** | **<0.001** |

*Dissolved Organic Carbon*

**Table S13. Differences between days, within treatments**

| Year | Treatment | Day1 - Day2 | Day1 - Day3 | Day2 - Day3 | Day3 - Day4 | Day4 - Day5 | Day5 - Day6 | Day6 - Day7 |
|---|---|---|---|---|---|---|---|---|
| 2022 | Control | 1 | **0.006** | **0.059** | 1 | 0.822 | **0.002** | 0.171 |
| | N | NA | 0.599 | NA | **0.006** | 0.997 | **<0.001** | **0.007** |
| | P | 1 | 1 | 1 | **<0.001** | **<0.001** | 0.999 | 0.066 |
| | NP | NA | 0.074 | NA | **0.001** | **0.026** | 0.08 | 0.324 |
| 2023 | Control | 1 | 0.293 | 0.9 | 0.992 | 0.997 | 0.529 | 0.204 |
| | N | 0.666 | 0.926 | 0.231 | 0.883 | **0.017** | 0.96 | 0.467 |
| | P | 1 | 0.931 | 0.945 | 0.747 | 1 | 0.998 | NA |
| | NP | 0.952 | 0.91 | 0.595 | 0.68 | 1 | 0.996 | 0.095 |
| | NPSi | NA | 0.906 | NA | 1 | **<0.001** | 1 | 0.96 |

**Table S14. Differences between treatments, within days**

| Year | Day | Control - N | Control - P | Control - NP | N - P | N - NP | NP - P | Control - NPSi | N - NPSi | NPSi - P | NP - NPSi |
|---|---|---|---|---|---|---|---|---|---|---|---|
| 2022 | 1 | 1 | 1 | 1 | 1 | 1 | 1 | | | | |
| | 2 | NA | 0.942 | NA | NA | NA | NA | | | | |
| | 3 | 0.116 | **0.003** | 0.71 | 0.519 | 0.62 | 0.059 | | | | |
| | 4 | 0.28 | **<0.001** | **0.004** | 0.123 | 0.29 | 0.966 | | | | |
| | 5 | 0.795 | **0.001** | 0.749 | **<0.001** | 0.233 | **0.021** | | | | |
| | 6 | 0.927 | 0.864 | **<0.001** | 0.998 | **<0.001** | **<0.001** | | | | |
| | 7 | 0.28 | 0.617 | **<0.001** | 0.934 | **0.002** | **<0.001** | | | | |
| 2023 | 1 | 0.566 | 0.566 | 0.566 | 1 | 1 | 1 | 0.652 | 1 | 1 | 1 |
| | 2 | 0.92 | 0.967 | 0.999 | 0.694 | 0.962 | 0.928 | NA | NA | NA | NA |
| | 3 | 0.988 | 0.987 | 0.989 | 1 | 1 | 1 | 0.997 | 1 | 1 | 1 |
| | 4 | 0.751 | 0.67 | 0.709 | 0.127 | 0.115 | 1 | 1 | 0.755 | 0.666 | 0.704 |
| | 5 | 0.644 | 0.891 | 0.959 | 0.99 | 0.956 | 0.999 | **0.001** | 0.062 | **0.019** | **0.01** |
| | 6 | 0.811 | 0.998 | 0.994 | 0.936 | 0.958 | 1 | 0.113 | **0.007** | 0.056 | **0.045** |
| | 7 | **0.039** | NA | **<0.001** | NA | 0.472 | NA | **0.006** | 0.787 | NA | 0.981 |

*Dissolved Inorganic Carbon*

**Table S15. Differences between days**

| Year | Treatment | Day1 - Day2 | Day1 - Day4 | Day1 - Day6 | Day1 - Day7 | Day2 - Day4 | Day2 - Day6 | Day2 - Day7 | Day4 - Day6 | Day4 - Day7 | Day6 - Day7 |
|---|---|---|---|---|---|---|---|---|---|---|---|
| 2022 | Control | 1 | 0.984 | 1 | 1 | 0.996 | 0.998 | 1 | 0.962 | 0.997 | 0.998 |
| | N | 0.998 | 0.997 | 1 | 0.189 | 1 | 0.984 | 0.324 | 0.982 | 0.334 | 0.127 |
| | P | 0.988 | 0.998 | 1 | 0.983 | 0.937 | 0.993 | 1 | 0.996 | 0.921 | 0.989 |
| | NP | 0.998 | 0.384 | 0.873 | 1 | 0.233 | 0.71 | 0.996 | 0.91 | 0.417 | 0.896 |
| Year | Treatment | Day1 - Day3 | Day1 - Day4 | Day1 - Day5 | Day1 - Day7 | Day3 - Day4 | Day3 - Day5 | Day3 - Day7 | Day4 - Day5 | Day4 - Day7 | Day5 - Day7 |
| 2023 | Control | 0.992 | 0.636 | 0.532 | 1 | 0.892 | 0.822 | 0.991 | 1 | 0.63 | 0.525 |
| | N | 1 | 0.768 | 0.627 | 0.999 | 0.785 | 0.671 | 0.999 | 1 | 0.87 | 0.798 |
| | P | 0.999 | 0.888 | 0.452 | 0.878 | 0.8 | 0.364 | 0.787 | 0.94 | 1 | 0.947 |
| | NP | 0.894 | 0.342 | **<0.001** | **0.001** | 0.099 | **<0.001** | **<0.001** | **<0.001** | 0.133 | **0.018** |
| | NPSi | 0.925 | **0.017** | **<0.001** | **<0.001** | 0.141 | **<0.001** | **<0.001** | **<0.001** | **<0.001** | **<0.001** |

**Table S16 Differences between treatments**

| Year | Day | Control - N | Control - P | Control - NP | N - P | N - NP | NP - P | Control - NPSi | N - NPSi | NPSi - P | NP - NPSi |
|---|---|---|---|---|---|---|---|---|---|---|---|
| 2022 | 1 | 1 | 1 | 1 | 1 | 1 | 1 | | | | |
| | 2 | 0.999 | 0.991 | 0.999 | 0.999 | 1 | 0.999 | | | | |
| | 4 | 0.998 | 0.871 | 0.147 | 0.936 | 0.205 | 0.495 | | | | |
| | 6 | 1 | 0.998 | 0.871 | 0.994 | 0.897 | 0.778 | | | | |
| | 7 | 0.235 | 0.987 | 0.996 | 0.394 | 0.157 | 0.945 | | | | |
| 2023 | 1 | 1 | 1 | 1 | 1 | 1 | 1 | 1 | 1 | 1 | 1 |
| | 3 | 0.996 | 0.976 | 0.797 | 0.999 | 0.945 | 0.984 | 0.998 | 0.957 | 0.892 | 0.613 |
| | 4 | 1 | 0.994 | 0.993 | 0.993 | 0.999 | 0.918 | 0.453 | 0.694 | 0.249 | 0.714 |
| | 5 | 1 | 1 | **< 0.001** | 0.999 | **<0.001** | **<0.001** | **<0.001** | **<0.001** | **<0.001** | **<0.001** |
| | 7 | 0.999 | 0.905 | **0.002** | 0.975 | **0.01** | **0.039** | **<0.001** | **<0.001** | **<0.001** | **0.002** |

*Total Alkalinity*

**Table S17. Differences between days**

| Year | Treatment | Day1 - Day2 | Day1 - Day4 | Day1 - Day6 | Day1 - Day7 | Day2 - Day4 | Day2 - Day6 | Day2 - Day7 | Day4 - Day6 | Day4 - Day7 | Day6 - Day7 |
|---|---|---|---|---|---|---|---|---|---|---|---|
| 2022 | Control | 0.906 | **0.015** | 0.626 | 0.779 | 0.113 | 0.982 | 0.999 | 0.302 | 0.194 | 0.999 |
| | P | 0.588 | 0.87 | 0.869 | 0.59 | 0.986 | 0.986 | 1 | 1 | 0.986 | 0.986 |
| | N | 0.788 | 0.087 | 0.261 | 0.096 | 0.567 | 0.883 | 0.595 | 0.978 | 1 | 0.984 |
| | NP | 0.864 | **0.053** | 0.082 | **0.031** | 0.346 | 0.454 | 0.238 | 1 | 0.999 | 0.993 |
| Year | Treatment | Day1 - Day3 | Day1 - Day4 | Day1 - Day5 | Day1 - Day7 | Day3 - Day4 | Day3 - Day5 | Day3 - Day7 | Day4 - Day5 | Day4 - Day7 | Day5 - Day7 |
| 2023 | Control | 1 | 0.898 | 0.961 | 0.979 | 0.891 | 0.954 | 0.994 | 1 | 0.641 | 0.77 |
| | N | 0.999 | 0.974 | 0.97 | 0.995 | 0.936 | 0.919 | 0.971 | 1 | 0.998 | 1 |
| | P | 0.999 | 0.967 | 0.804 | 0.969 | 0.91 | 0.696 | 0.913 | 0.992 | 1 | 0.991 |
| | NP | 0.991 | 0.99 | 0.147 | 0.99 | 1 | 0.095 | 1 | 0.093 | 1 | 0.094 |
| | NPSi | **<0.001** | **<0.001** | **<0.001** | **<0.001** | 0.872 | **0.008** | 1 | **<0.001** | 0.944 | **0.005** |

**Table S18. Differences between treatments**

| Year | Day | Control - N | Control - P | Control - NP | N - P | N - NP | NP - P | Control - NPSi | N - NPSi | NPSi - P | NP - NPSi |
|---|---|---|---|---|---|---|---|---|---|---|---|
| 2022 | 1 | 1 | 1 | 1 | 1 | 1 | 1 | | | | |
| | 2 | 0.994 | 0.931 | 1 | 0.986 | 0.999 | 0.96 | | | | |
| | 4 | 0.879 | 0.09 | 0.954 | 0.349 | 0.996 | 0.245 | | | | |
| | 6 | 0.914 | 0.97 | 0.595 | 0.692 | 0.93 | 0.336 | | | | |
| | 7 | 0.482 | 0.989 | 0.227 | 0.677 | 0.958 | 0.376 | | | | |
| 2023 | 1 | 1 | 1 | 1 | 1 | 1 | 1 | 1 | 1 | 1 | 1 |
| | 3 | 1 | 1 | 0.998 | 1 | 1 | 1 | **<0.001** | **<0.001** | **<0.001** | **<0.001** |
| | 4 | 1 | 0.999 | 0.762 | 1 | 0.902 | 0.876 | **0.024** | 0.082 | **0.013** | **0.001** |
| | 5 | 1 | 0.996 | 0.562 | 0.993 | 0.533 | 0.782 | **<0.001** | **<0.001** | **<0.001** | **<0.001** |
| | 7 | 0.91 | 0.821 | 1 | 1 | 0.944 | 0.881 | **<0.001** | **0.001** | **0.002** | **<0.001** |
